# Origin of diversity in spatial social dilemmas

**DOI:** 10.1101/2021.03.21.436334

**Authors:** Christoph Hauert, Michael Doebeli

## Abstract

Cooperative investments in social dilemmas can spontaneously diversify into stably co-existing high and low contributors in well-mixed populations. Here we extend the analysis to emerging diversity in (spatially) structured populations. Using pair approximation we derive analytical expressions for the invasion fitness of rare mutants in structured populations, which then yields a spatial adaptive dynamics framework. This allows us to predict changes arising from population structures in terms of existence and location of singular strategies, as well as their convergence and evolutionary stability as compared to well-mixed populations. Based on spatial adaptive dynamics and extensive individual based simulations, we find that spatial structure has significant and varied impacts on evolutionary diversification in continuous social dilemmas. More specifically, spatial adaptive dynamics suggests that spontaneous diversification through evolutionary branching is suppressed, but simulations show that spatial dimensions offer new modes of diversification that are driven by an interplay of finite-size mutations and population structures. Even though spatial adaptive dynamics is unable to capture these new modes, they can still be understood based on an invasion analysis. In particular, population structures alter invasion fitness and can open up new regions in trait space where mutants can invade, but that may not be accessible to small mutational steps. Instead, stochastically appearing larger mutations or sequences of smaller mutations in a particular direction are required to bridge regions of unfavourable traits. The net effect is that spatial structure tends to promote diversification, especially when selection is strong.

## 1 Introduction

Social dilemmas (Dawes, 1980; Hauert et al., 2006) are important mathematical metaphors for studying the problem of cooperation. A famous example is the tragedy of the commons (Hardin, 1968), which results in overexploitation of communal resources. The best studied models of social dilemmas are the prisoner’s dilemma and the snowdrift game (Doebeli and Hauert, 2005). Traditionally, such models are often restricted to the two distinct strategies of cooperate, *C*, and defect, *D*. However, these models can easily be extended to describe a continuous range of cooperative investments into a public good. In such continuous games, the aim is to study the evolutionary dynamics of the level of investment.

For example, in the donation game (Sigmund, 2010), which is the most prominent version of the prisoner’s dilemma, cooperators confer a benefit *b* to their interaction partners at a cost *c* to themselves. Defection entails neither costs nor confers benefits. With *b > c >* 0, both individuals prefer mutual cooperation, which pays *b − c*, over a zero payoff for mutual defection. However, the temptation to shirk costs and free ride on benefits provided by the partner undermines cooperation to the detriment of all. In fact, defection is the dominant strategy because it results in higher payoffs regardless of the partner’s strategy. This conflict of interest between the individual and the group represents the hallmark of social dilemmas.

A natural translation of the donation game to continuous traits is based on cost and benefit functions, *C*(*x*) and *B*(*x*), where the trait *x ∈* [0, *x*_max_] represents a level of cooperative investment that can vary continuously. In the spirit of the donation game, an individual with strategy *x* interacting with an *y*-strategist obtains a payoff *P*(*x, y*) = *B*(*y*) *− C*(*x*). Assuming that (*i*) zero investments incur no costs and provide no benefits, *B*(0) = *C*(0) = 0, (*ii*) benefits exceed costs, *B*(*x*) *> C*(*x*) ≥ 0, and (*iii*) are increasing functions, *B*′(*x*), *C*′(*x*) ≥ 0, recovers the social dilemma of the donation game for continuous investment levels: the level of investment invariably evolves to zero, which corresponds to pure defection, despite the fact that both players would be better off at non-zero investment levels. The reason is that an actor can only influence the cost *C*(*x*) of an interaction, but not the received benefit *B*(*y*), and hence selection can only act to minimize costs.

In a weaker form of a social dilemma, the snowdrift game (Hauert and Doebeli, 2004; Sugden, 1986), cooperators also provide benefits, *b*, at a cost, *c*. However, benefits are now accessible to both individuals and accumulate in a discounted manner. For example, yeast secretes enzymes for extra cellular digestion of sucrose (Greig and Travisano, 2004). While access to nutrients is crucial, the marginal value of additional resources diminishes and may exceed the intake capacity (Hauert et al., 2006). As a result, the social dilemma remains in effect, because the most favourable outcome of mutual cooperation remains prone to cheating, but the dilemma is relaxed because it pays to cooperate against defectors. Again, it is straightforward to formulate a continuous version of the snowdrift game. In contrast to the donation game, in the continuous snowdrift game the benefits depend on the strategies of both interacting individuals. For example, for simplicity one can assume that benefits depend on the aggregate investment levels of both players, so that the benefit is a function *B*(*x* + *y*). Assuming a cost function *C*(*x*), an *x* strategist interacting with strategy *y* obtains a payoff *P*(*x, y*) = *B*(*x* + *y*) *− C*(*x*), again with *B*(0) = *C*(0) = 0, *B*(*x*) *> C*(*x*) ≥ 0 and *B*′(*x*), *C*′(*x*) ≥ 0, as before, but with the additional constraint *C*(*x*) *> B*(*x* + *y*) *− B*(*y*) for sufficiently large *y* so that the increased return from investments of an individual do not outweigh its costs when interacting with high investors. Typically this is achieved by saturating benefit functions, *B*′′(*x*) *<* 0.

The gradual evolution of continuous traits can be described using the framework of adaptive dynamics (Dieckmann and Law, 1996; Geritz et al., 1997, 1998; Metz et al., 1996). The central quantity is the invasion fitness *f*(*x, y*), which is the growth rate of a rare mutant *y* in a monomorphic resident population with trait *x*. To derive the adaptive dynamics of the trait *x*, one considers the selection gradient *D*(*x*) = *∂*_*y*_*f*(*x, y*)|_*y*=*x*_. If *D*(*x*) *>* 0, nearby mutants with trait values *y > x* can invade, whereas for *D*(*x*) *<* 0 nearby traits with *y < x* invade. For example, it is easy to see that in the continuous snowdrift game, *D*(0) *>* 0 if *B*′(*x*) *> C*′(*x*) ≥ 0, so that in contrast to the continuous donation game, the pure defector strategy *x* = 0 is not evolutionarily stable (Doebeli et al., 2004). Traits *x*^*\**^ with *D*(*x*^*\**^) = 0 are called singular traits and are convergent stable if the Jacobian of the selection gradient, 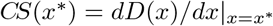, is negative, whereas evolutionary stability requires the Hessian of the fitness, 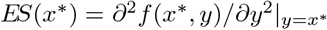, to be negative, i.e. that the invasion fitness *f*(*x, y*) has a (local) maximum at *x*^*\**^. Since the conditions for convergent stability and for evolutionary stability are different, in general, a singular trait *x*^*\**^ may be convergent stable but not evolutionarily stable. This results in the intriguing dynamical feature of evolutionary branching, where an initially monomorphic population evolutionarily splits into two (or more) co-existing traits (Doebeli, 2011).

For social dilemmas with a discrete set of strategies, there exists extensive literature on the effects of spatial structures on the dynamics of the social dilemma (Allen et al., 2017; Débarre et al., 2014; Nowak and May, 1992; Ohtsuki et al., 2006; Szabó and Fáth, 2007). In particular, it has been shown in many incarnations that spatial structure can promote cooperative strategies, and hence that spatial structure can have a profound effect on game dynamics. However, far less is known about the effects of spatial structure on continuous games (Allen et al., 2013; Le Gaillard et al., 2003), and on evolutionary diversification in particular. In well-mixed populations, evolutionary branching has been shown to occur generically in the continuous snowdrift game (Doebeli et al., 2004), in which cooperative investments can diversify into stably co-existing high and low contributors. Similar observations have been made for the “tragedy of the commune”, a game that is “dual” to the tragedy of the commons in the sense that the trait *x* represents the amount of consumption of a public good. In this case, diversification results in a coexisting coalition of unequal consumers of a public good (Killingback et al., 2010).

Here we study evolutionary diversification in continuous spatial games. To derive the corresponding spatial adaptive dynamics, we first use the technique of pair approximation (Matsuda et al., 1992; Szabó and Fáth, 2007; van Baalen, 2000) to describe the frequency dynamics of pairs of resident and mutant traits in order to obtain an analytical expression for the invasion fitness of a rare mutant in a structured resident population. We then use the invasion fitness function to determine singular trait values and their convergence and evolutionary stability. Analytical approximations and predictions are tested against individual-based simulations of the corresponding spatial games. It turns out that in structured populations evolutionary branching is suppressed but easily compensated for by other modes of diversification especially for stronger selection such that evolutionary diversification represents a robust feature of continuous spatial games.

## 2 Spatial invasion fitness and adaptive dynamics

The dynamics of spatial games is generally complex (Ibsen-Jensen et al., 2015; Szabó and Hauert, 2002) and depends on the details of the updating process (Débarre et al., 2014; Ohtsuki et al., 2006). In particular, the sequence of birth and death events can be of crucial importance in spatially structured populations. For example, in the classical donation game with selection on birth rates in spatially structured populations, cooperators invariably disappear for birth-death updating but are able to thrive for death-birth updating (provided that the benefits exceed the *k*-fold costs, *b > ck* (Ohtsuki et al., 2006)). Here we analyze the spatial evolutionary dynamics for the more interesting case of death-birth updating for social dilemmas with continuous traits, as exemplified by the continuous prisoner’s dilemma and the continuous snowdrift game. For birth-death updating the effects of population structures are less pronounced, and the corresponding results are relegated to SI Text S4.

In the following, we assume that the total population size is constant, and that spatially structured populations are represented by lattices in which each site is occupied by one individual. Each individual interacts with a limited number of local neighbours, and we assume this number, *k*, to be the same for all individuals. We first consider a case where there are two types of players in the structured population: a mutant type with trait value *y*, and a resident type with trait value *x* (where *x* and *y* denote investment strategies in a continuous game). If the mutant has *j* other mutants among its *k* neighbours, the mutant payoff is *π*_*m*_(*j*) = [(*k−j*)*P*(*y, x*)+*jP*(*y, y*)]*/k*. Similarly, the payoff of a resident with *j* mutant neighbours is given by *π*_*r*_(*j*) = [*jP*(*x, y*) +(*k − j*)*P*(*x, x*)]*/k*. The payoffs, *π*_*m*_(*j*), *π*_*r*_(*j*), of mutants and residents from interactions with their *k* neighbours determines the birth rates as *b*_*m*_(*j*) = exp(*wπ*_*m*_(*j*)) and *b*_*r*_(*j*) = exp(*wπ*_*r*_(*j*)), where *w >* 0 denotes the strength of selection. The birth rate is proportional to the probability of taking over an empty site for which a given mutant or resident individual competes. For *w ≪* 1 selection is weak and differences in payoffs result in minor differences in birthrates, and hence in small differences in probabilities of winning competition for an empty site. With strong selection, *w ≫* 1, payoff differences are amplified in the corresponding birthrates. This exponential payoff-to-birthrate map has several convenient features: (*i*) ensures positive birthrates, (*ii*) admits easy conversion to probabilities for reproduction, (*iii*) selection can be arbitrarily strong, (*iv*) for weak selection the more traditional form of birthrates *b*_*i*_(*j*) *≈* 1 + *wπ*_*i*_(*j*), *i* = *m, r*, is recovered.

In well-mixed populations the current state is simply given by the frequency of mutants and residents, respectively. In contrast, in structured populations the state space is immense because it involves all possible configurations. Pair approximation offers a convenient framework to account for corrections arising from spatial arrangements (Matsuda et al., 1992; Szabó and Fáth, 2007; van Baalen, 2000). Instead of simply tracking the frequencies of mutants and residents, pair approximation considers the frequencies of neighbouring strategy pairs. We denote the frequencies of mutant-mutant, mutant-resident, resident-mutant and resident-resident pairs by *p*_*mm*_, *p*_*mr*_, *p*_*rm*_, and *p*_*rr*_, respectively. This reduces to two dynamical equations because *p*_*mr*_ = *p*_*rm*_ and *p*_*mm*_ + *p*_*mr*_ + *p*_*rm*_ + *p*_*rr*_ = 1 must hold. The most informative quantities are the global mutant frequency, *p*_*m*_ = *p*_*mm*_ + *p*_*mr*_, and the local mutant density *q*_*m*|*m*_ = *p*_*mm*_*/p*_*m*_, i.e. the conditional probability that a neighbour of a mutant is also a mutant. Note that for rare mutants, *p*_*m*_ *«* 1, their local densities need not be small. The derivation of the corresponding dynamical equations depends on the details of the microscopic updating.

The death-birth process with selection on birth in structured populations results in local competition: an individual is selected to die uniformly randomly from the whole population, then its *k* neighbours compete to repopulate the newly vacated site. They succeed with a probability proportional to their birthrates. Note that payoffs, and hence birth rates, are based on interactions with all neighbours, including the neighbour that may subsequently be chosen to die (uniformly at random) and its vacant site subject to recolonization by the offspring of one of its neighbours. To determine the dynamics of *p*_*m*_ and *q*_*m*|*m*_, we first note that *p*_*m*_ only changes when a resident is replaced by a mutant, or when a mutant is replaced by a resident. We denote by 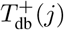 the probability that a resident dies that has *j* mutant neighbours, and that the removed resident is replaced by one of its mutant neighbours. Similarly, we denote by 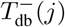 the probability that a mutant dies that has *j* mutant neighbours, and that the removed mutant is replaced by one of its resident neighbours.

The dynamical equations for the pair approximation with death-birth updating are then given by:

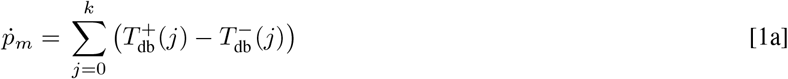

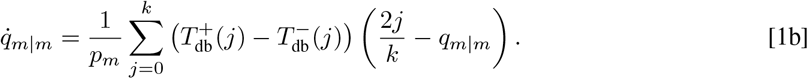

Here the difference between the rates of increase, 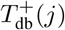, and decrease, 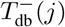 mutant numbers is determined by:

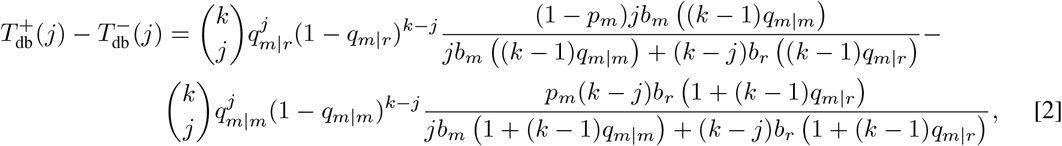

where *b*_*m*_(*v*) and *b*_*r*_(*v*) denote the birth rates of mutants and residents, respectively, with an average number of *v* mutants in their neighbourhood (see SI Text S1 for details). The dynamical system given by Eq. 1 is an approximation for the frequency dynamics of any given resident-mutant pair.

To obtain the invasion fitness, we note that the dynamics of spatial invasion unfolds in two stages: mutants quickly establish a local (pseudo) equilibrium and then gradually increase (or decrease) in frequency (Fu et al., 2010; Langer et al., 2008; Le Gaillard et al., 2003). Formally, this is reflected in a time scale separation in the limit of rare mutants, *p*_*m*_ *«* 1, between the slow global frequency dynamics, Eq. 1a, and the fast local pair density, Eq. 1b. As a consequence, to calculate invasion fitness we assume that the local pair density of the mutant is at the equilibrium 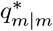 determined by Eq. 1b, i.e. by 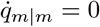. As shown in SI Text S1, the invasion fitness of mutants, *f*(*x, y*), defined as their per capita growth rate, 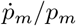, in the limit *p*_*m*_ *→* 0, then becomes:

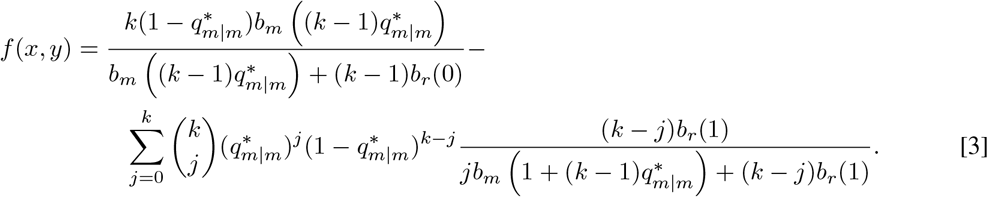

Even though the solution to 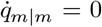 is analytically inaccessible, in general, the equilibrium 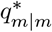 can be approximated using a Taylor expansion if |*y − x*| *«* 1 (see SI Text S1):

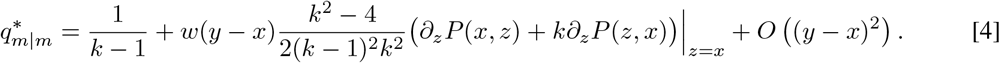

It follows that in the limit *y → x*, mutants with at least one resident neighbour have, on average, one mutant neighbour among their *k −* 1 other neighbours. Note, mutants with no resident neighbours are uninteresting because they are unable to initiate a change in the population configuration. Interestingly, this limit of the local pair configuration is fairly robust with respect to changes in the updating process (c.f. Eq. S24 in SI Text S4 for birth-death updating). Moreover, in this limit a rare mutation with positive invasion fitness is guaranteed to eventually take over (see (Dieckmann and Law, 1996) and SI Text S1).

Using Eqs. 3 and (4) the selection gradient, 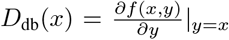, as well as its Jabobian, 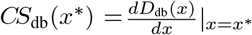, and the Hessian of fitness, 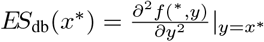, at a singular point *x\** can be calculated as:

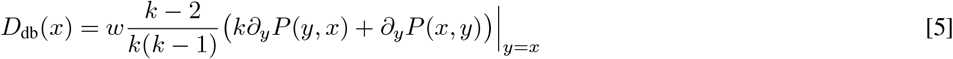

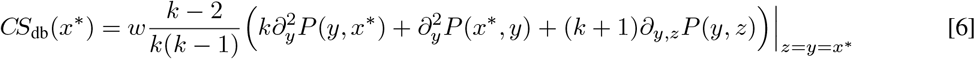

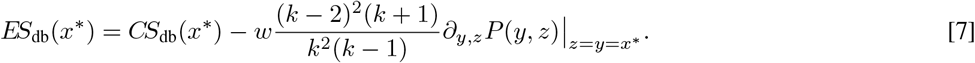

## 3 Continuous social dilemmas in space

We now apply the approach described above to specific continuous spatial games. To get a sense of the predictive power of the analytical approximations, we compare the analytical approximations to results from corresponding individual-based models.

### 3.1 Continuous prisoner’s dilemma

In the continuous prisoner’s dilemma the payoff to an individual with strategy *y* interacting with an *x*-strategist is *P*(*y, x*) = *B*(*x*) *− C*(*y*). This implies that the selection gradient in well-mixed populations is proportional to *−C*′(*x*), and hence, assuming monotonously increasing costs, no singular strategy exists apart from the pure defection state *x* = 0, and the population always evolves to that state, regardless of how large the benefits of cooperation are. In contrast, for death-birth updating in structured populations the selection gradient, Eq. 5, becomes

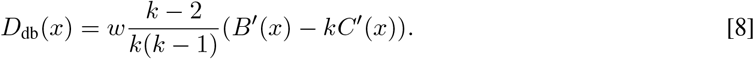

Thus, in structured populations a singular strategy, *x*^*\**^, may exist as a solution to *D*_db_(*x*^*\**^) = 0. If *x*^*\**^ exists and is convergent stable then it is also evolutionarily stable because the two stability conditions are identical (the mixed derivatives on the right hand side of Eqs. 6 and (7) are zero). In particular, cooperation can be maintained in spatially structured populations, a result that is of course in line with classical theory (Nowak and May, 1992). More specifically, cooperative investments can increase if the marginal benefits, *B*′(*x*), exceed the *k*-fold marginal costs, *C*′(*x*), which is reminiscent of the *b > ck*-rule in the traditional donation game (Ohtsuki et al., 2006).

#### 3.1.1 Linear costs and benefits

The evolutionary analysis becomes particularly simple for linear benefit and cost functions *B*(*x*) = *x* and *C*(*x*) = *rx*, where *r* denotes the (marginal) cost-to-benefit ratio. Again, population structures are capable of supporting cooperation for sufficiently small *r*. More precisely, the selection gradient 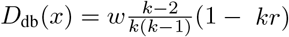 is reduced to a constant (c.f. Eq. 8), and hence no singular strategy exists. However, the gradient changes sign for *r*^*\**^ = 1*/k*. Thus, for favourable cost-to-benefit ratios, *r < r*^*\**^, investments increase to their maximum, whereas for harsher conditions, *r > r*^*\**^ cooperative investments cannot be sustained, see Fig. 1.

**Fig. 1.**
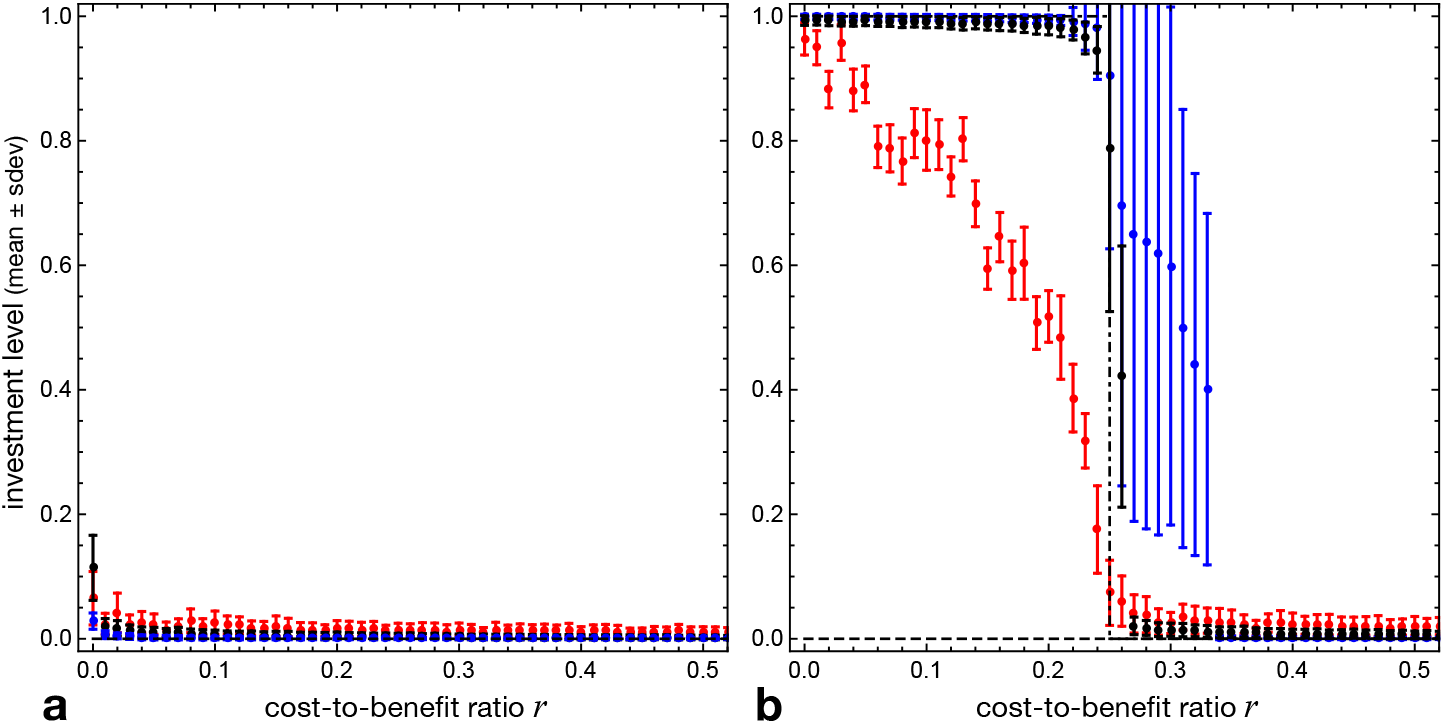
Equilibrium investment levels (mean *±* standard deviation) in individual-based simulations of the linear continuous prisoner’s dilemma (*B*(*x*) = *x, C*(*x*) = *rx*) as a function of the cost-to-benefit ratio, *r*, for weak selection, *w* = 1 (red), moderate, *w* = 10 (black), and strong selection, *w* = 100 (blue). **a** in well-mixed populations with *N* = 10^4^ individuals cooperative investments are close to zero regardless of *r* as predicted by adaptive dynamics (dashed line). The small variance further decreases for stronger selection emphasizing the disadvantage of mutants with higher investments. **b** for populations with *k* = 4 neighbours spatial adaptive dynamics predicts a threshold *r*^*\**^ = 1*/k* (dash-dotted-line) below which investments reach the maximum but disappear above. Simulations on a 100 *×* 100 lattice confirm the trend but reveal an interesting susceptibility to noise: for weak selection (red) the maximum investment is not reached; intermediate selection (black) essentially follows pair approximation, while strong selection (blue) maintains non-zero investment levels beyond the predicted threshold. The large variation suggests co-existence of different traits and is confirmed through sample snapshots of the spatial configuration in Fig. 2i (click here for interactive online simulations).

Interestingly, these analytical predictions are not always borne out in individual-based models when the selection strength, *w*, is large. The reasons for the differences can be appreciated by looking at pairwise invasibility plots (PIP), which show the regions in (*x, y*)-space in which a mutant *y* can invade a resident *x*, i.e., regions for which the invasion fitness *f*(*x, y*) is positive. To construct these plots, we first solve the dynamical equation 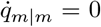, Eq. 1b, to find the equilibrium 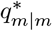 for given resident and mutant trait values *x, y* and then determine the sign of the corresponding invasion fitness, Eq. 3. An example is shown in Fig. 2g with *r >* 1*/k* so that adaptive dynamics predicts evolution to pure defection. For mutants *y* sufficiently close to the resident *x, f*(*x, y*) *>* 0 only holds for *y < x*, so that evolution by (infinitesimally) small mutations, as envisaged in adaptive dynamics, indeed leads to ever smaller investment. However, the PIP also shows positive invasion fitness for mutants *y > x* if *y* is sufficiently larger than *x* (white area above the diagonal in Fig. 2g). The width of the region of unfavourable mutants decreases with increasing selection strength *w*, i.e. the gap becomes easier to bridge (but does not depend on the resident trait *x*; grey area in Fig. 2g for *w* = 50 and black area for *w* = 100). In individual-based models, mutations are normally distributed around the parental trait and of finite size such that stochastically appearing larger mutations or sequences of smaller mutations in the right direction can give rise to the co-existence of high and low investors (see Fig. 2i). Thus, evolutionary diversification is possible in individual-based models despite the absence of evolutionary branching points in the adaptive dynamics, and even if the analytical prediction is evolution of pure defection. Note that this mode of evolutionary diversification is a feature of spatial structure that does not occur in the corresponding well-mixed models.

**Fig. 2.**
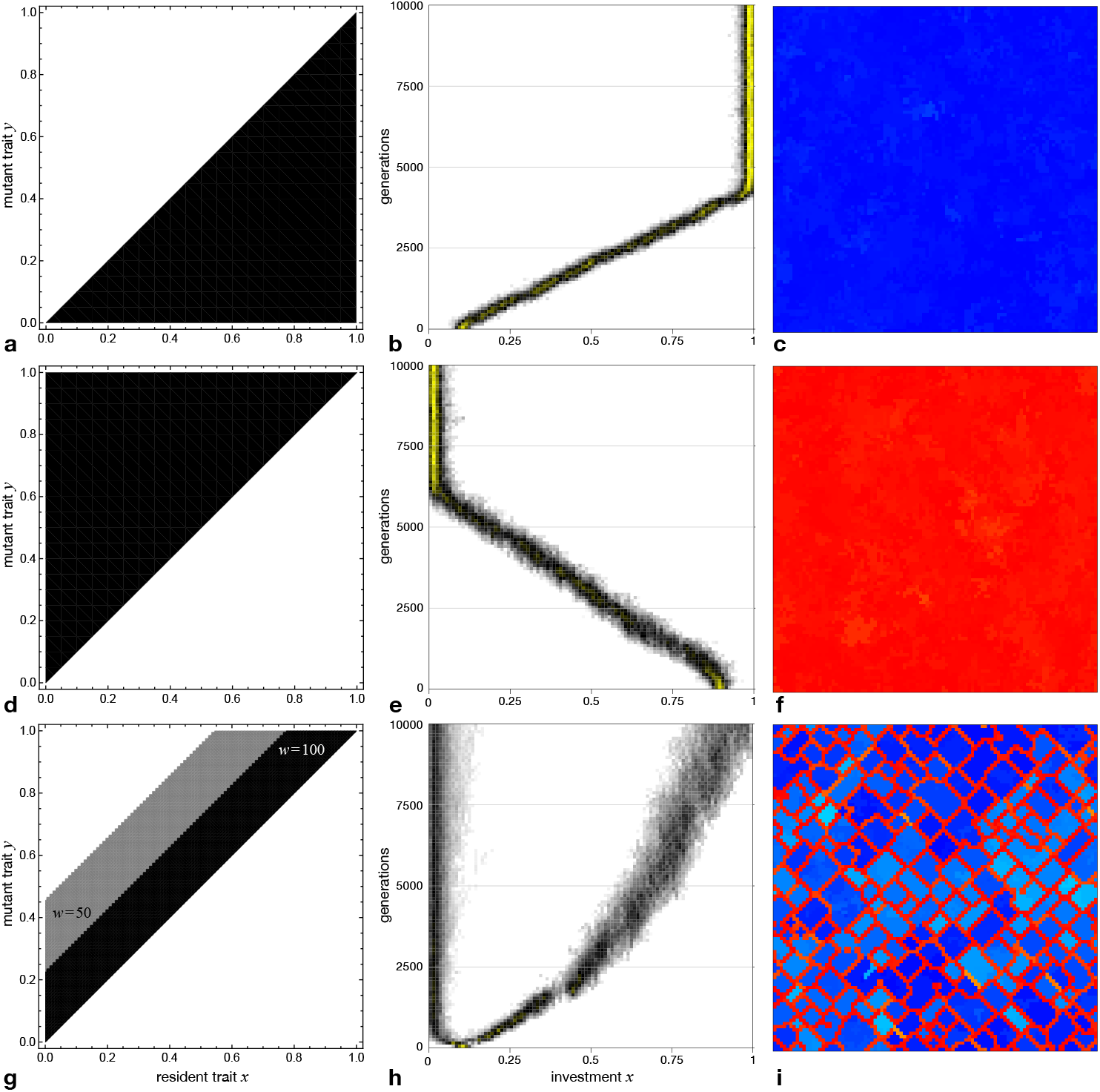
Linear continuous prisoner’s dilemma in 100 *×* 100 lattice populations with *k* = 4 neighbours. Evolutionary dynamics is illustrated for favourable cost-to-benefit ratios, *r* = 0.1 *<* 1*/k* (**a**-**c**, click here for interactive online simulations, (Hauert, 2021)), and for harsher conditions, *r* = 0.3 *>* 1*/k*, with moderate selection, *w* = 10 (**d**-**f**, interactive simulations) as well as strong selection, *w* = 100 (**g**-**i** interactive simulations). The left column shows the pairwise invasibility plots (PIP), which indicate whether mutant traits are capable of invading a particular resident population (white regions) or not (black regions). The middle column shows the evolutionary trajectory of the distribution of investments over time in individual-based simulations (darker shades indicate higher trait densities in the population with the highest densities in yellow). The right column depicts snapshots of the population configuration at the end of the simulation runs. The colour hue indicates the investment levels ranging from low (red) to intermediate (green) and high (blue). For *r <* 1*/k* higher investors can always invade and eventually the maximum investment is reached (**a**-**c**), regardless of selection strength. The situation is reversed for *r >* 1*/k* and weak to moderate selection where only lower investors can invade and investments dwindle to zero (**d**-**f**). Interestingly, for strong selection in lattice populations not only lower investors can invade for *r >* 1*/k* but also those that invest significantly more than the resident (**g**-**f**). Nevertheless adaptive dynamics predicts that investments evolve to zero because of the assumption that mutations are small, which restrict the dynamics to the diagonal of the PIP. However, individual-based simulations show that rare larger mutations or a sequence of smaller ones can give rise to the co-existence of high and low investors. The outcome does not depend on the initial configuration of the population.

##### Saturating benefits

The continuous prisoner’s dilemma was first introduced in Killingback et al. (1999) with saturating benefits *B*(*x*) = *b*_0_ (1 *−* exp(*−b*_1_*x*)) and linear costs *C*(*x*) = *c*_0_*x* (*b*_0_, *b*_1_, *c*_0_ *>* 0). Using individual-based simulations these authors showed that structured populations are capable of supporting cooperation, whereas well-mixed populations are not. Spatial adaptive dynamics provides an analytical perspective on these results. For the given cost and benefit functions, we have:

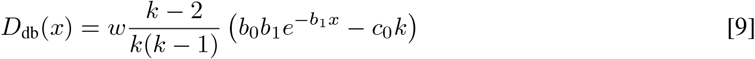

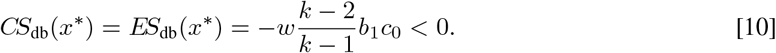

There is a singular point at

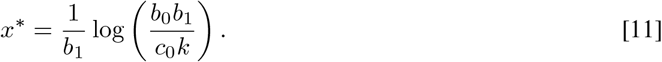

The singular point is always convergent stable as well as evolutionarily stable (see Fig. 3b). Interestingly, the adaptive dynamics analysis again misses subtle but intriguing effects arising from strong selection. A pairwise invasibility plot for strong selection is shown in Fig. 3d) and reveals that *x*^*\**^ is susceptible to invasion by mutants with slightly higher investments. In individual-based models, this effectively turns *x*^*\**^ into a (degenerate) branching point, i.e., a starting point for evolutionary diversification. The diversification into coexisting high and low investors has already been observed in Killingback et al. (1999), but the underlying mechanism had not been addressed. The earlier results were based on a different, deterministic update rule, according to which a focal individual imitated the strategy of the best performing neighbour, including itself, but this update rule essentially corresponds to death-birth updating with very strong selection (the only difference being that the focal individual is removed). Hence the diversification reported in Killingback et al. (1999) is of the same type as the one seen here for strong selection.

**Fig. 3.**
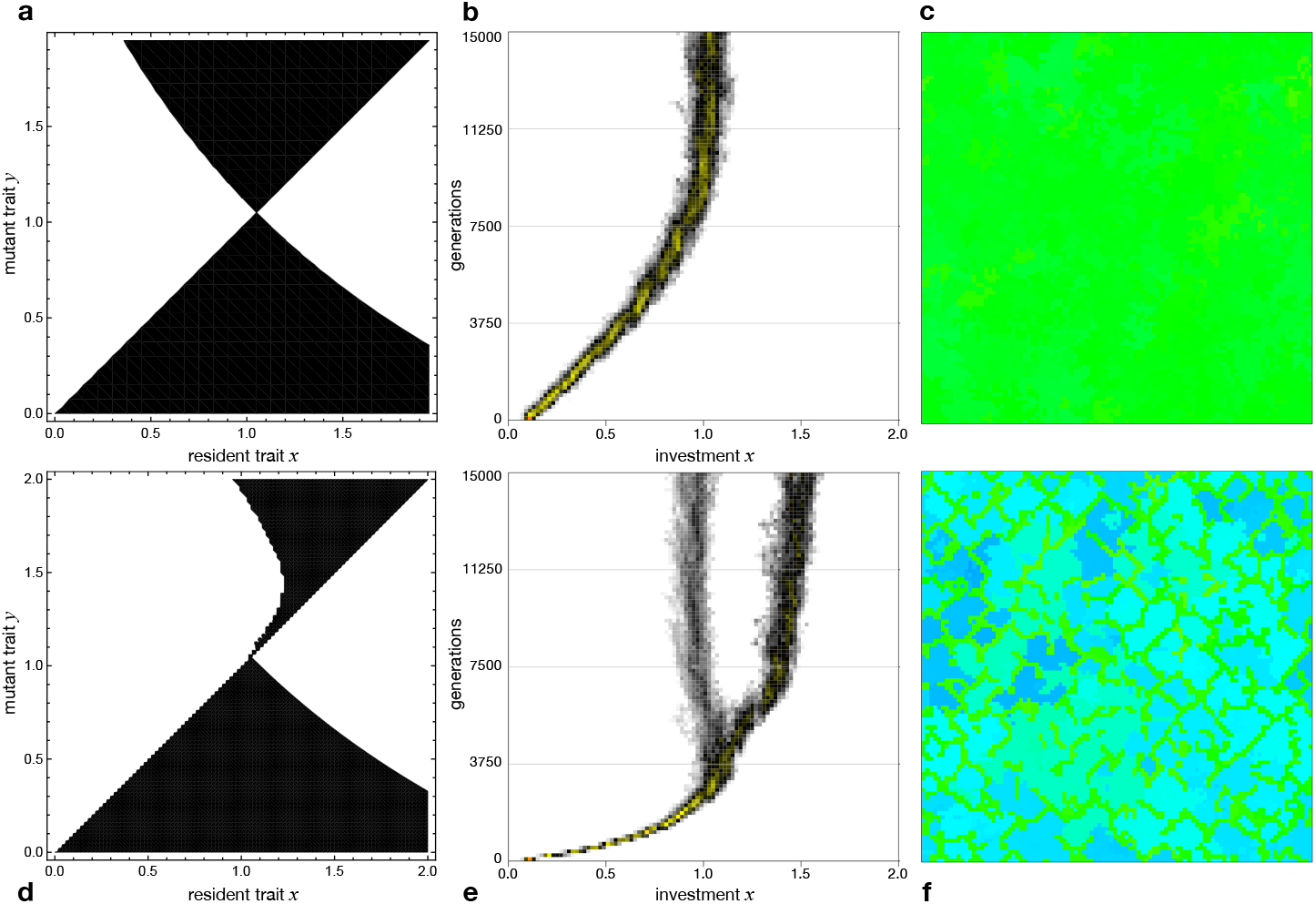
Continuous prisoner’s dilemma with saturating benefits *B*(*x*) = *b*_0_ (1 *−* exp(*−b*_1_*x*)) and linear costs *C*(*x*) = *c*_0_*x* for *b*_0_ = 8, *b*_1_ = 1, *c*_0_ = 0.7 in 100 *×* 100 lattice populations for weak selection, *w* = 1 (top row, click here for interactive online simulations, (Hauert, 2021)) and strong selection, *w* = 100 (bottom row, interactive simulations). The pairwise invasibility plots (PIP, left column) show that higher investing mutants can invade for low resident investments and lower investing mutants can invaded high investing residents. However, near the singular investment level, 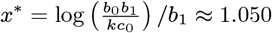, selection strength gives rise to interesting differences in the dynamics. **a** for weaker selection, *w* = 1, *x*^*\**^ is evolutionarily stable. This is confirmed through individual-based simulations. **B** depicts the investment distribution over time (darker shades indicate higher trait densities in the population with the highest densities in yellow). **c** shows a snapshot of the spatial configuration at the end of the simulation. The colour hue indicates the investment level ranging from low (red) to intermediate (green) to high (blue). In contrast, **d** for strong selection, *w* = 100, *x*^*\**^ can be invaded but only by higher investing mutants. As a consequence, a degenerate form of evolutionary branching may occur. Individual-based simulations confirm branching already for *w* = 10 (**e, f**, interactive simulations).

### 3.2 Continuous snowdrift game

In the continuous snowdrift game the payoff to an individual with strategy *y* interacting with an *x*-strategist is *P*(*y, x*) = *B*(*x* + *y*) *− C*(*y*). Note that this generalization to continuous strategies does not imply that every interaction between an individual with strategy *x* and another with strategy *y* invariably results in a snowdrift game. In fact, snowdrift games in this narrower sense appear only if the two traits *x, y* straddle the singular strategy *x*^*\**^. Depending on the cost and benefit functions, as well as on the strategies *x* and *y*, any kind of 2 *×* 2 game can arise, including a prisoner’s dilemma, in which the lower investing strategy dominates (Doebeli et al., 2004, 2013). In this sense the framework of the continuous snowdrift game actually encompasses the gist of the continuous prisoner’s dilemma for sufficiently high costs – just as the classical snowdrift game turns into a prisoner’s dilemma (Hauert and Doebeli, 2004).

The selection gradient for the continuous snowdrift game in structured populations, Eq. 5, is given by

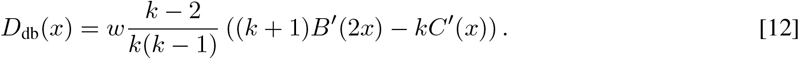

Thus, singular strategies *x*^*\**^ may exist, and if *x*^*\**^ exists, it is convergent stable if

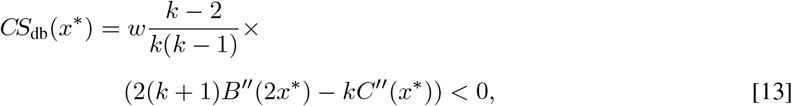

and evolutionarily stable if

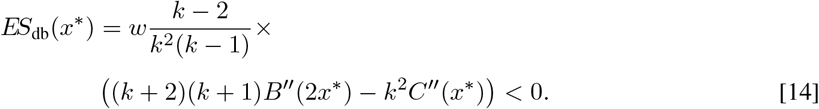

In particular, the conditions for convergent and evolutionary stability are different, which indicates the potential for evolutionary branching and hence the evolutionary emergence and co-existence of high investing cooperators and low investing defectors. The above conditions are the spatial analogs of the analysis for well-mixed continuous snowdrift games reported in (Doebeli et al., 2004). Also note that for linear cost and benefit functions it is not possible to satisfy all constraints for the continuous snowdrift game and hence the simplest case is given by quadratic costs and benefits.

#### 3.2.1 Quadratic costs and benefits

For suitable cost and benefit functions the (spatial) adaptive dynamics is analytically accessible. Here we focus on the quadratic cost and benefit functions used in (Doebeli et al., 2004) for the well-mixed case: *B*(*x*) = *b*_1_*x* + *b*_2_*x*^2^ and *C*(*x*) = *c*_1_*x* + *c*_2_*x*^2^. We assume that the evolving trait is confined to the interval [0, 1]. The benefit and cost functions satisfy the criteria *B*(0) = *C*(0) = 0, *B*(*x*), *C*(*x*) ≥ 0 and *B*′′(*x*) *<* 0 for *x ∈* [0, 1] provided that *b*_1_, *c*_1_ *>* 0, *b*_2_, *c*_2_ *<* 0 as well as *b*_1_ *< −*4*b*_2_ and *c*_1_ *< −*2*c*_2_. The first two conditions ensure that costs and benefits are increasing yet saturating, while the latter two ensure that *B*(2*x*) and *C*(*x*) are increasing over the entire trait interval, i.e. their respective maximum occurs for *x* ≥ 1.

For small resident values *x*, the selection gradient Eq. 12 becomes 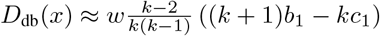, and hence cooperative investments evolve away from zero provided that 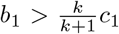, which is slightly less restrictive than the corresponding condition *b*_1_ *> c*_1_ in the well-mixed case (Doebeli et al., 2004).

For the above quadratic costs and benefits the singular strategy is given by

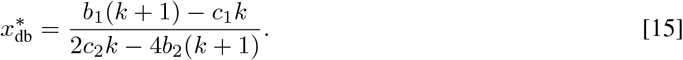

The numerator of Eq. 15 is positive if and only if the condition for evolution away from zero, 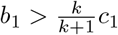, is satisfied. Similarly, the denominator of Eq. 15 is positive if and only if 2*b*_2_(*k* + 1) *< c*_2_*k* (recall *b*_2_, *c*_2_ *<* 0). It is clear from Eq. 13 that this is also the condition for convergence stability. Thus, if a singular point, *x*^*\**^, exists, cooperative traits either evolve away from zero and *x*^*\**^ is convergent stable, or the trait cannot evolve away from zero and *x*^*\**^ is a repellor.

We note that if the singular point exists, *x*^*\**^, it is shifted towards smaller investments for a given set of parameters, as compared to well-mixed populations with the same parameters: 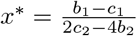 (Doebeli et al., 2004). Furthermore, the condition for convergence stability, 2*b*_2_(*k* + 1) *< c*_2_*k*, is less restrictive than the corresponding condition 2*b*_2_ *< c*_2_ in the well-mixed case. Finally, from Eq. 14 we see that the condition for evolutionary stability is *b*_2_(*k* + 2)(*k* + 1) *< c*_2_*k*^2^, which is again less restrictive than the corresponding condition *b*_2_ *< c*_2_ in the well-mixed case. Combining the two stability conditions shows that evolutionary branching occurs for 2*b*_2_(*k* + 1)*/k < c*_2_ *< b*_2_(*k* + 1)(*k* + 2)*/k*^2^. Thus, the analytical approach based on pair approximation suggests that population structures tend to inhibit evolutionary diversification by decreasing the range of parameters for which the singular point is an evolutionary branching point (c.f. Fig. 4a & c).

In order to explore the evolutionary dynamics more specifically and for a range of parameters, it is convenient to fix two parameters, say *b*_2_ and *c*_1_, which then leaves a range for the other two parameters (subject to the constraints listed above, i.e. 0 *< b*_1_ *< −*4*b*_2_ and 0 *> c*_2_ *> −c*_1_*/*2). Strictly speaking, *b*_1_ *< c*_1_ violates one of the assumptions of the continuous snowdrift game at least for small *x*, namely that *B*(*x*) *> C*(*x*). It means that for small *x* lower investors dominate and investments are expected to dwindle and disappear, which actually recovers the continuous prisoner’s dilemma scenario and cooperative investments cannot take off because *D*(*x*) *<* 0. In this sense the parameter space depicted in Fig. 4 encompasses the gist of the continuous analogs of both the prisoner’s dilemma and the snowdrift game.

**Fig. 4.**
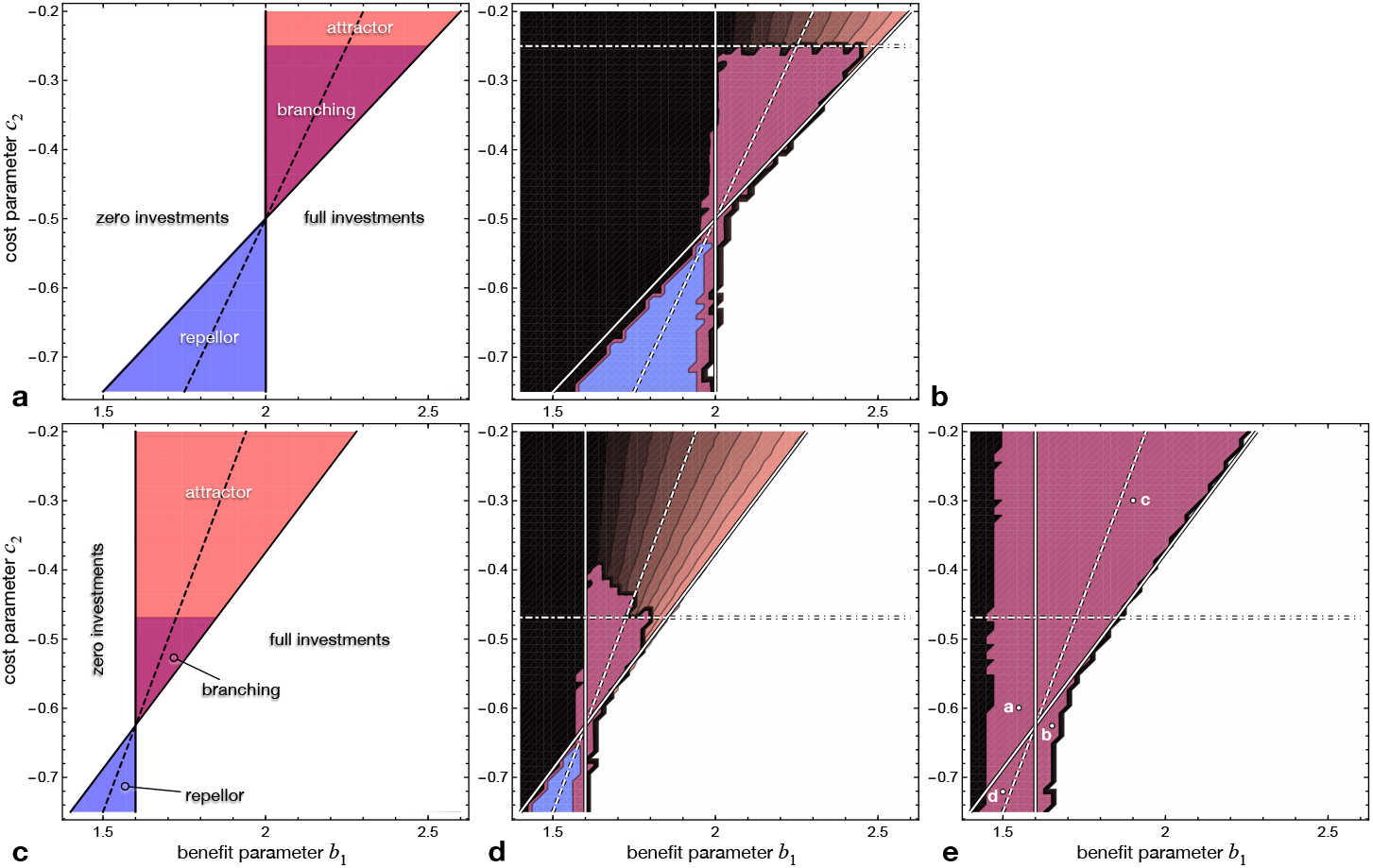
Continuous snowdrift game with quadratic benefit and cost functions, *B*(*x*) = *b*_1_*x* + *b*_2_*x*^2^ and *C*(*x*) = *c*_1_*x* + *c*_2_*x*^2^. Evolutionary outcomes are shown as a function of the benefit parameter *b*_1_ and cost parameter *c*_2_ with *b*_2_ = *−*1*/*4 and *c*_1_ = 2. Note that *b*_1_ *<* 2 violates the assumption *B*(*x*) *> C*(*x*) at least for small *x* and hence effectively mimics the characteristics of the prisoner’s dilemma. **a** analytical predictions based on adaptive dynamics in well-mixed populations and **b** results from individual-based simulations for populations with *N* = 10^4^ individuals.\ **c** analytical predictions based on spatial adaptive dynamics and complementing individual-based simulations on 100 *×* 100 lattices for **d** moderate selection, *w* = 10 (click here for interactive online simulations), and **e** strong selection. In lattice populations the parameter region admitting singular strategies is shifted to both smaller values of *b*_1_ and of *c*_2_ and the size of the region admitting evolutionary branching is markedly smaller than in well-mixed populations (**a, c**). Interestingly, spatial adaptive dynamics predicts branching only for parameters where defection dominates in well-mixed populations, *b*_1_ *< c*_1_, mimicking the continuous prisoner’s dilemma. For weak to moderate selection predictions by adaptive dynamics (**a, c**) are in good agreement with results from individual-based simulations (**b, d**), where equilibrium investment levels range from the minimum (black) to intermediate (grey) and the maximum (white) augmented by convergent stability (red) and evolutionary instability (blue) with the overlapping region indicating evolutionary branching (maroon) in adaptive dynamics and diversification in simulations. For strong selection (**e**) striking differences arise with a much increased region of diversification. The points labelled *a-d* indicate the parameter settings for the invasion analysis in Fig. 5. Note that the automated classification of investment distributions becomes more difficult whenever the singular investment *x*^*\**^ is close to zero or one (for details see SI Text S2).

This results in five salient regimes for the evolutionary dynamics (Doebeli et al., 2004): if *x*^*\**^ does not exist, investments evolve either to their (*i*) maximum or (*ii*) minimum, depending on the sign of the selection gradient. The latter case recovers the essence of the continuous prisoner’s dilemma and arises if *b*_1_ *< c*_1_ (at least for small *x*). If *x*^*\**^ exists but is not convergent stable, it is (*iii*) a repellor, and the evolutionary outcome depends on the initial investment, *x*_0_, such that for *x*_0_ *> x*^*\**^ investments evolve to the maximum, and for *x*_0_ *< x*^*\**^ investments evolve to zero. Conversely, if *x*^*\**^ is both convergent stable and evolutionarily stable it is (*iv*) an attractor of stable intermediate investments, which represent the evolutionary end state. Finally, if *x*^*\**^ is convergent stable but not evolutionarily stable, it is (*v*) an evolutionary branching point and hence a potential starting point for evolutionary diversification into co-existing high and low investors.

Figure 4 compares the different dynamical regimes for well-mixed and structured populations in terms of the parameters *b*_1_ and *c*_2_ based on analytical predictions derived from adaptive dynamics in well-mixed populations and for structured populations based on pair approximation as well as individual-based simulations (see SI Text S2 for simulation details). Interestingly, due to the shift of the singular point *x*^*\**^ towards smaller investments as compared to well-mixed populations, branching now only occurs for parameters that result in negative selection gradients in well-mixed populations (i.e. drive investments to zero, at least for small *x*) and hence qualitatively mimic the prisoner’s dilemma.

For weaker selection strengths, predictions based on (spatial) adaptive dynamics and individual-based simulations turn out to be in surprisingly good agreement given that the effects of space are reduced to mere pair correlations, Fig. 4b & d, while significant differences arise for stronger selection, Fig. 4e. In particular, the range of parameters leading to diversification is increased in individual-based models compared to the predictions based on pair approximation. This effect is particularly pronounced when selection is strong (see Fig. 4e). With strong selection, diversification expands into parameter regions that result in monomorphic traits at minimal, maximal or stable intermediate investments, as well as into regions of bi-stability (i.e., with *x*^*\**^ a repellor) for weaker selection.

Qualitatively, these discrepancies may not be that surprising, given that pair approximation cannot account for larger-scale spatial structures, such such as large clusters of cooperators, or frozen, filamentous regions of defectors (see Fig. 2i). Again the main reason for the discrepancies are finite-size mutations and non-monomorphic populations in individual-based models, which differ from the assumptions for adaptive dynamics. With finite mutations and finite population variances, unfavourable regions in the invasion fitness landscape can more easily be crossed by mutational steps to reach regions in which coexistence is feasible. This can be seen by using pairwise invasibility plots, PIP, to depict regions of mutual invasibility between pairs of trait values. In essence this means that population structures open new avenues for evolutionary diversification where divergent selection is not solely based on evolutionary branching.

Figure 5 shows PIPs for four different regimes where spatial adaptive dynamics does not predict branching but diversification emerges in individual-based simulations on lattices (c.f. Fig. 4e with each parameter set marked by the letters a, b, c, and d, respectively). In Fig. 5a the selection gradient is negative everywhere, and in Fig. 5b it is positive everywhere (indicated by the white regions below and above the diagonal, respectively). In both cases there is no singular point. However, for small values of the resident in Fig. 5a (large values in Fig. 5b) mutants with sufficiently larger (smaller) investment levels than the resident can invade and coexist with the resident. Regions of coexistence are characterized by the fact that mutants can invade residents and vice versa. Graphically this can be captured by *mutual* invasibility plots, PIP^2^, see Fig. 5e, f. Moreover, for persistent coexistence evolution must drive divergence of the coexisting branches (as indicated by the vector field in Fig. 5e, f). For strong selection, the impermissible regions shrink (grey and black areas in Fig. 5). In individual-based models with sufficiently strong selection, coexisting trait pairs eventually occur due to finite-size mutations. Divergence then results in persistent evolutionary diversification into distinct trait groups of high and low investors.

**Fig. 5.**
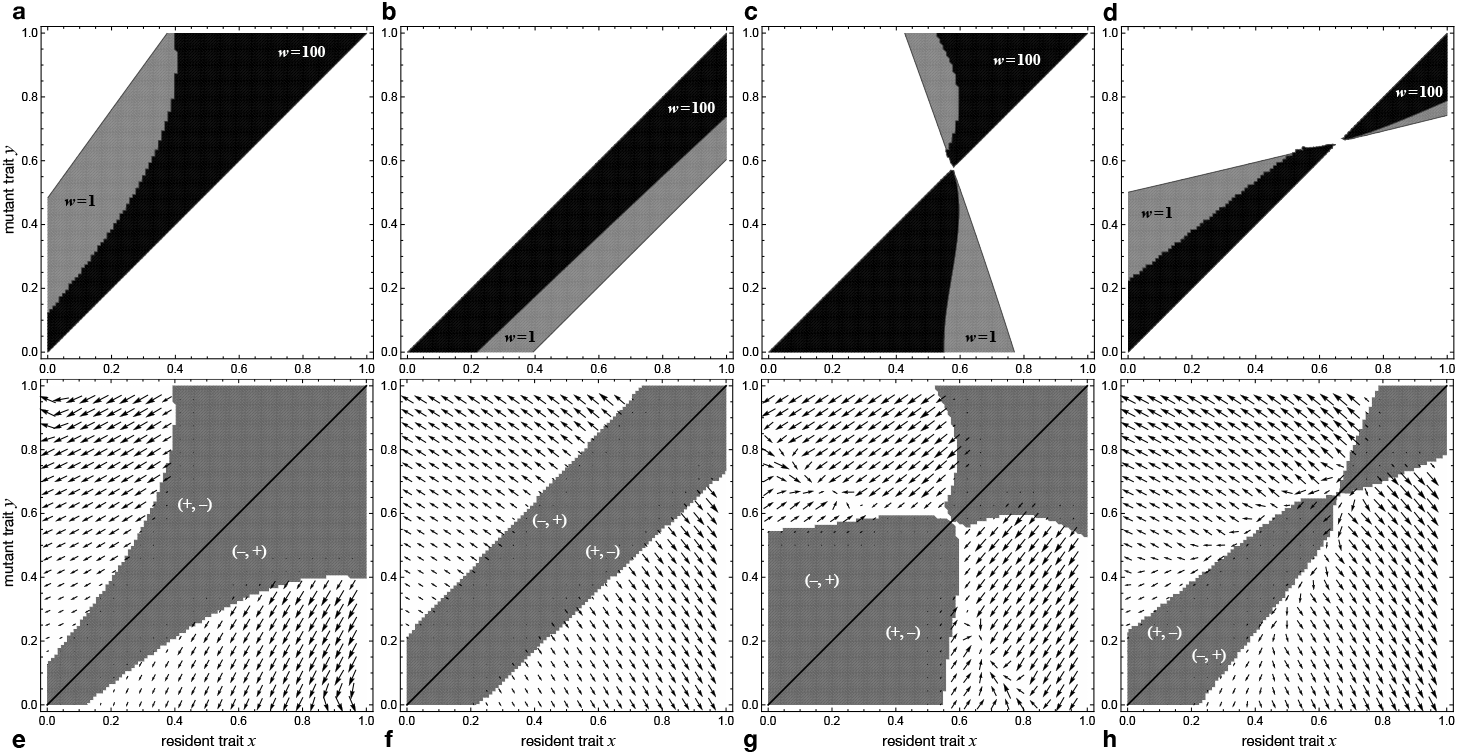
Spatial modes of diversification in the continuous snowdrift game with quadratic benefit and cost functions. **a**-**d** depict pairwise invasibility plots (PIP, top row) for the four scenarios illustrating increased spatial diversification due to strong selection (c.f. parameter combinations *a-d* marked in Fig. 4). In all cases the width of the region of disadvantageous mutants decreases with selection strength (grey for *w* = 1; black for *w* = 100). **a** the PIP suggests gradual evolution towards minimal investments, except for smaller resident traits, where not only lower investing mutants can invade but also those making markedly higher investing (click here for interactive online simulations). **b** higher investing mutants can always invade, but so can traits investing markedly less (interactive simulations). **C** selection strength distorts the PIP in the vicinity of the convergent and evolutionarily stable *x*^*\**^ resulting in a degenerate form of branching (c.f. Fig. 3, interactive simulations). **d** the PIP indicates that *x*^*\**^ is a repellor such that residents with *x < x*^*\**^ are invaded by lower investors while those with *x > x*^*\**^ by higher investors. However, as a consequence of strong selection, mutants with markedly higher (lower) investments can also invade (interactive simulations). **e**-**h** depict corresponding plots of regions of *mutual* invasibility (PIP^2^, white regions). Regions where mutants or residents are unable to invade (grey) are marked with (+, *−*) and (*−*, +), respectively. The vector field shows the divergence (see SI Text S3 for details) and indicates the direction of selection for two co-existing residents based on analytical approximations of the spatial invasion dynamics. In all cases divergence drives the traits away from the diagonal and hence preserves diversity. Parameters: *b*_2_ = *−*1*/*4, *c*_1_ = 2, **a** *b*_1_ = 1.55, *c*_2_ = *−*0.6; **b** *b*_1_ = 1.65, *c*_2_ = *−*0.625; **c** *b*_1_ = 1.9, *c*_2_ = *−*0.3; **d** *b*_1_ = 1.5, *c*_2_ = *−*0.72;

In Fig. 5c & d, singular points exist and are evolutionarily stable or a repellor, respectively. In both case many mutant-resident pairings in the vicinity of the singular point can co-exist. As a result, evolutionary stability is marginal in the former case and in the latter case the region of impermissible mutants around the diagonal is narrow. The effect is again more pronounced for strong selection. In either case finite-size mutations in individual-based models can easily lead to coexistence of diverging phenotypic branches (see Fig. 5g, h), and hence result in evolutionary diversification.

## 4 Discussion

Spontaneous diversification in social dilemmas into high and low investors tends to be facilitated by population structures, especially when selection is strong. However, at the same time, classical evolutionary branching tends to be inhibited, but compensated for by other modes of diversification. Here we derive an extension of adaptive dynamics for continuous games in (spatially) structured populations based on pair approximation, which tracks the frequencies of mutant-resident pairs during invasion. It turns out that predictions derived from this spatial adaptive dynamics framework are independent of selection strength. More precisely, selection strength only scales the magnitude of the selection gradient as well as of convergent and evolutionary stability but neither affects the location of singular strategies nor their stability. Nevertheless, from the invasion analysis of mutant traits *y* into resident populations *x*, as well as from individual-based simulations, it is clear that selection strength has a significant impact on the dynamics in general, and on diversification in particular. During the invasion process the local configuration probability of mutant pairs *q*_*m*|*m*_ is of crucial importance and changes with selection strength, see Eq. 4. However, in the adaptive dynamics limit *y → x* the pair configuration probability reduces to a constant *q*_*m*|*m*_ *→* 1*/*(*k −* 1). Even though this is the same value as obtained in the limit of weak selection, *w →* 0, it is important to note that the limit *y → x* does not necessarily imply weak selection. Thus, the different modes of diversification cannot be understood based on spatial adaptive dynamics alone but require a more comprehensive analysis of invasion dynamics.

Structured populations offer new modes of diversification that are driven by an interplay of finite-size mutations and population structures. Trait variation is more easily maintained in structured populations due to the slower spreading of advantageous traits as compared to well-mixed populations. Spatial adaptive dynamics is unable to capture these new modes of diversification because of the underlying assumption that the resident population is composed of discrete traits (monomorphic before branching and dimorphic or polymorphic after branching). Nevertheless, invasion analysis and pairwise invasibility plots, in particular, provide an intuitive interpretation for additional pathways to diversification that are introduced by spatial structures and further promoted for increasing selection strength. Even in the vicinity of evolutionarily stable investments, trait combinations exist that admit mutual invasion and hence can coexist. However, such states cannot evolve through adaptive dynamics but are nevertheless accessible to trait distributions around the evolutionarily stable trait and can drive a degenerate form of branching. Moreover, spatial invasion fitness can open up new regions in trait space for mutant invasion. However, those regions need not be accessible by small mutational steps, and instead require stochastically appearing larger mutations or sequences of smaller mutations that allow to bridge regions of unfavourable traits.

Previous attempts at amalgamating adaptive dynamics and spatial structure have not observed spontaneous diversification or evolutionary branching. More specifically, Allen et al. (2013) augment adaptive dynamics by structure coefficients (Tarnita et al., 2011), which imply weak selection. Even though they do not report evolutionary branching, their approach aligns well with our results. In particular, one of their examples follows Killingback et al. (1999) and corresponds to our section 3.1.2 and Fig. 3, and refers to a system which indeed exhibits spontaneous diversification through a degenerate form of branching – but only for stronger selection. Their other example considers a multi-dimensional case where the payoffs linearly depend on trait values. As a consequence evolutionary branching is not expected because the difference between convergent and evolutionary stability depends on second derivatives, and hence the two criteria necessarily coincide in their models. However, for the continuous snowdrift game their setup should equally predict evolutionary branching for appropriate parameters. Another attempt at spatial adaptive dynamics considers spatial structure in the form of demes (Parvinen et al., 2017). However, quite naturally such deme structures suppress branching because in the long run it is impossible to maintain multiple co-existing traits in small demes that quickly become homogeneous. Instead, branching is only possible if demes are large enough to support branching in every single one.

Interestingly, in well-mixed populations evolutionary branching is only observed for the continuous snowdrift game, where two distinct traits of high and low investors can co-exist and essentially engage in a classical (discrete) snowdrift game. In contrast, in structured populations with death-birth updating, evolutionary branching is only observed for prisoner’s dilemma type interactions where lower investments invariably dominate higher ones, which applies both in the continuous prisoner’s dilemma as well as the continuous snowdrift game with sufficiently high costs. The reason for this surprising difference can be understood intuitively by considering the preferred spatial configurations in the two classical (discrete) games: in the prisoner’s dilemma cooperators form compact clusters to reduce exploitation by defectors (minimize surface), while in the snowdrift game filament like clusters form because it is advantageous to adopt a strategy that is different from that of the interaction partners (maximize surface). In the continuous variants of those games it is naturally much harder to maintain and spread distinct traits in fragmented filament-like structures because they are more prone to effects of noise than compact clusters. Effectively this fragmentation inhibits evolutionary branching because diverging traits tend to trigger further fragmentation and as a consequence do not survive long enough to get established and form their own branch. In contrast, the compact clusters promoted by the prisoner’s dilemma provide structural protection for higher investors and thus help drive diversification.

Because of global competition the spatial dynamics for birth-death updating is (unsurprisingly) much closer to results for well-mixed populations. For example, evolutionary branching was again only observed for continuous snowdrift game. Also because of global competition, structured populations are updated in a non-uniform manner. In particular, regions of high payoffs experience a much higher turnover than regions of low payoffs. For strong selection this can result in almost frozen parts of the population. As a consequence unsuccessful traits are able to stay around for long times and, in some cases, those traits turn out to be advantageous again at later times when the surroundings have sufficiently changed, so that the stragglers then contribute to diversification. This mode of diversification, however, introduces historical contingencies where the evolutionary end state can sensitively depend on the initial configuration.

Overall, we find that evolutionary diversification is a robust feature of continuous spatial games, and that spatial structure can sometimes hinder, but generally promotes diversification through modes of diversification that complement traditional evolutionary branching.

## Acknowledgements

This research was funded by the National Science and Engineering Research Council of Canada (NSERC) Discovery grants RGPIN-2015-05795 to CH and 219930 to MD.

## Data availability

The source code for the individual-based simulations is available at https://github.com/evoludolab/IBS-DSCG.

## Supporting Information

### S1 Spatial adaptive dynamics: death-birth updating

For continuous games in spatially structured populations, the frequency dynamics of two types *x* and *y* can be modelled using pair approximation. This reduces the complex spatial dynamics to four pair configuration probabilities: *p*_*mm*_, *p*_*mr*_, *p*_*rm*_ and *p*_*rr*_, which indicate the frequencies of mutant-mutant, mutant-resident, resident-mutant and resident-resident pairs, respectively, with *p*_*mm*_ + *p*_*mr*_ + *p*_*rm*_ + *p*_*rr*_ = 1. The frequency of mutants is given by *p*_*m*_ = *p*_*mm*_ + *p*_*mr*_ and that of residents by *p*_*r*_ = *p*_*rr*_ + *p*_*rm*_. For consistency, the frequency of mutants can also be expressed as *p*_*m*_ = *p*_*mm*_ + *p*_*rm*_ by considering links that point towards mutants rather than away from them. Consequently *p*_*rm*_ = *p*_*mr*_ must hold and the dynamics is reduced to two dynamical variables. The most intuitive pair is given by the global frequency of mutants, *p*_*m*_, and their local pair density as represented by the conditional probability, *q*_*m*|*m*_ = *p*_*mm*_*/p*_*m*_. The respective rates of change depend on the microscopic updating procedure.

For death-birth updating, first the birthrate of all individuals is calculated by taking the payoffs from interactions with their *k* neighbours into account (we assume regular spatial structures in which each individual has the same number of neighbours *k*). An individual is then uniformly at random selected to die, and its *k* neighbours compete to repopulate the vacant site with their offspring. The probability of success is proportional to the birthrate. We note that this setup implies that competition occurs at the local scale, and that life expectancy is the same for all individuals. This is in contrast to birth-death updating where competition occurs globally and individuals with successful neighbours have a lower life expectancy (see SI Text S4).

The frequency of mutants and mutant-mutant pairs increases whenever a resident dies and a mutant neighbour repopulates the vacated site. For a resident with *j* mutant neighbours this happens with probability:

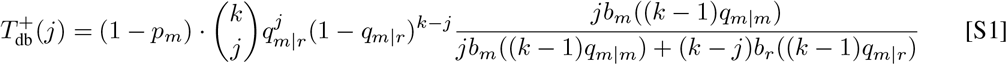

with *q*_*m*|*r*_ = (1 *− q*_*m*|*m*_)*p*_*m*_*/*(1 *− p*_*m*_), and where *b*_*r*_(*v*), *b*_*m*_(*v*) denote the birthrates of residents and mutants with an average number of *v* mutants among their neighbours. The first term in Eq. S1 indicates the probability that a resident dies, the second denotes the probability that it had *j* mutant neighbours and the last term is the probability that one of them reproduces. Similarly, the frequency of mutants and mutant-mutant pairs decreases if a mutant dies and one of its resident neighbours reproduces. For a mutant with *j* mutant neighbours this happens with probability:

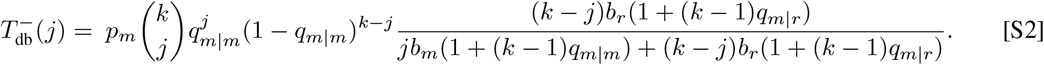

All other transitions do not alter the composition of the population, and hence the 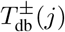 for *j* = 0,…, *k* define the rate of change of the frequency of mutants:

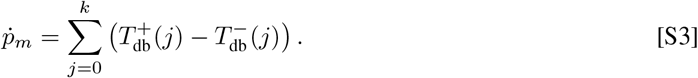

The derivation of the rate of change of *q*_*m*|*m*_ is a bit trickier and it helps to start with changes in *p*_*mm*_. First, consider a mutant that has successfully replaced a resident neighbour. This resident neighbour has one mutant neighbour (the reproducing individual) as well as an expected *q*_*m*|*r*_(*k −* 1) further mutants among its remaining *k −* 1 neighbours. Hence *p*_*mm*_ increases at a rate proportional to (1 + *q*_*m*|*r*_(*k −* 1))*/*(*Nk/*2), where *Nk/*2 indicates the normalization, i.e. the total number of (undirected) links in a population of size *N*. Similarly, if a resident has successfully replaced a mutant neighbour, *p*_*mm*_ decreases at a rate *q*_*m*|*m*_(*k −* 1)*/*(*Nk/*2), because the mutant neighbour has itself *q*_*m*|*m*_(*k −* 1) mutant neighbours (note, one is a resident with certainty, i.e. the reproducing individual). Thus the rate of change of *p*_*mm*_ becomes:

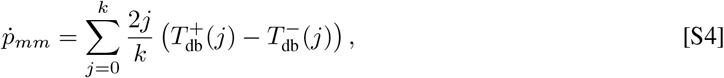

where the term 1*/N* has been omitted because the rates of change in both *p*_*m*_ and *p*_*mm*_ are proportional to the inverse population size and hence this factor can be absorbed through a constant rescaling of time. Finally, using 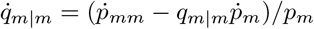 yields:

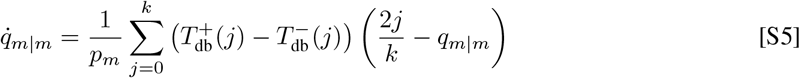

The dynamical equations for the global, Eq. S3, and local, Eq. S5, frequencies of mutants make up Eq. 1 in the main text.

The invasion fitness of a rare mutant trait *y* in a resident *x* is defined by 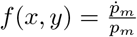 in the limit *p*_*m*_ *→* 0.

We first note that the leading order of the global dynamics, Eq. S3, is *O*(*p*_*m*_), while the local dynamics, Eq. S5, scales with *O*(1) and hence happens much faster when mutants are rare, *p*_*m*_ *«* 1. This results in a convenient separation of time scales, so that we can use the equilibrium 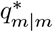 of Eq. S5 in Eq. S3 to calculate the invasion fitness. In the limit *p*_*m*_ *→* 0, the sum over 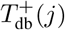 in Eq. S3 and divided by *p*_*m*_ reduces to:

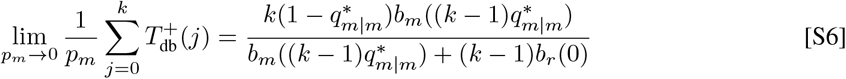

using 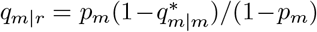). In contrast, the sum over 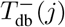 simplifies only marginally by cancelling the common factor *p*_*m*_ and *q*_*m*|*r*_ *→* 0. Thus the invasion fitness becomes:

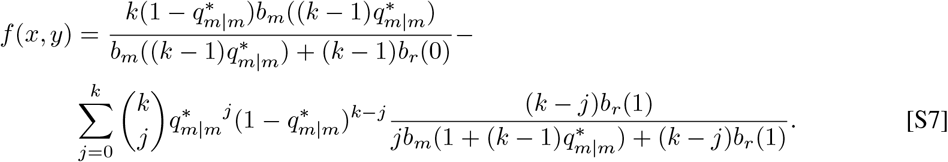

In order to calculate 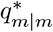, we first note that in the limit of rare mutants, *p*_*m*_ *→* 0, and for mutant traits *y* close to the resident trait *x*, Eq. S5 somewhat simplifies to

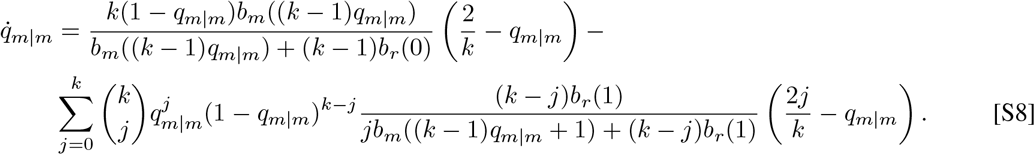

Second, a Taylor expansion of the right-hand-side of Eq. S8 in *y* around *x* yields, up to first order:

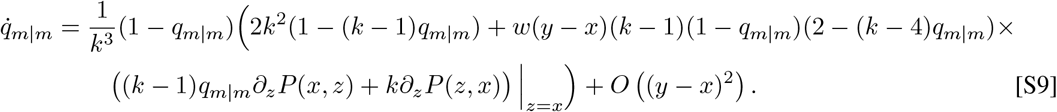

Thus, in this approximation, the equilibrium 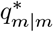 is given by the roots of a third order polynomial (plus the trivial, uninteresting root *q*_*m*|*m*_ = 1). To circumvent further analytical challenges, the zeroth order approximation of 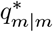 in *y* near *x* can be obtained by solving for the roots of Eq. S9 for *y* = *x*, which yields 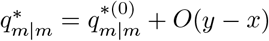 with 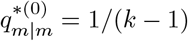.

Next, the first order approximation of 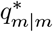 is obtained by implicit differentiation of Eq. S9 with respect to *y* (keeping in mind that *q*_*m*|*m*_ is a function of *y*) and evaluation at *y* = *x*. Setting the expression to zero yields an equation for the zeroth and first order coefficients, 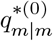 and 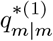, of the Taylor expansion at the equilibrium 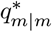 for *y* near *x*:

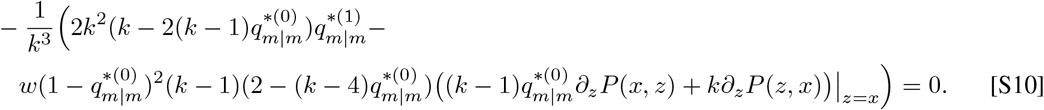

Solving for the first order coefficient 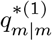 using 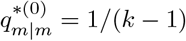 then yields:

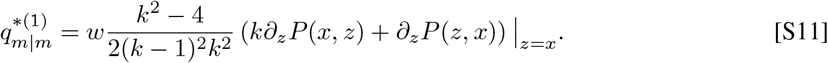

Assembling all the pieces finally results in the first order expansion in *y* of the local pair density equilibrium, 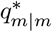, around *x*, see Eq. 4 in the main text. The upshot is that this allows to simplify the invasion fitness to a function of the mutant and resident traits only.

### S2 Individual-based simulations

Simulations were carried out for populations of size *N* = 10^4^ and run for 10^4^ generations. In well-mixed populations the payoffs and hence birthrates of individuals are based on interactions with other members of the population selected uniformly at random. In structured populations, individuals are arranged on a 100 *×* 100 lattice and interact with their *k* = 4 nearest neighbours, corresponding to the von Neumann neighbourhood.

The birthrate *b*_*i*_ = exp(*wπ*_*i*_) of every individual *i* is based on its payoff *π*_*i*_ from a single interaction with a random partner in well-mixed populations and on the average payoff of interactions with all *k* = 4 neighbours in lattice populations. The exponential mapping ensures that birthrates are always positive and *w* denotes the selection strength. For death-birth updating each computational update step consists of a death event followed by a birth event and vice versa for birth-death updating. This ensures that the population size remains constant.

For death-birth updating an individual is uniformly at random selected to die and then all neighbours compete for reproduction. With a probability proportional to their birthrates one neighbour succeeds in reproducing and placing an offspring on the vacated site. For birth-death updating an individual from the entire population is selected with a probability proportional to its birthrate. Its offspring replaces one of the parents neighbours selected uniformly at random. In well-mixed populations the neighbours include all *N −* 1 members of the population for death-birth updating (and *N* for birth-death, i.e. including the parent). On lattices competition is restricted to the *k* neighbours of the vacated site or the parent, respectively. Whenever an individual reproduces, the offspring inherits the parental strategy. However, with probability 0.01 a mutation occurs and the offspring’s strategy is drawn from a Gaussian distribution around the parental strategy with standard deviation *σ*_mut_ = 0.01.

The initial trait values are randomly assigned from a Gaussian distribution with mean *x*_0_ and standard deviation *σ*_0_ = 0.01. In most cases simulations start at low trait values of *x*_0_ = 0.1 because the most basic question is whether cooperative investments can evolve out of non-cooperative ancestral states. However, in some cases *x*_0_ = *x*_max_ *−* 0.1 is used to either demonstrate that not even high initial cooperative investments can prevent the demise of cooperation or that the evolutionary end state is robust with respect to the initial configuration. In particular, the latter does not apply in the case of a repellor and hence allows to discriminate between global attractors of the dynamics and bi-stable scenarios.

The classification of the evolutionary outcomes for the continuous snowdrift game that result in (*i*) diversification or (*ii*) bi-stability deserve some further remarks (see Fig. S1 and Fig. 4 in the main text): (*i*) diversification is based on the bi-modality coefficient

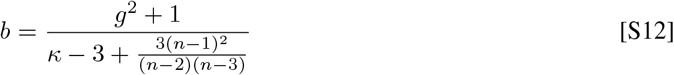

where *n* denotes the sample size, *κ* the kurtosis, and *g* the skewness of the trait distribution (Freeman and Dale, 2013). Uniform and exponential distributions have *b*_*u*_ = 5*/*9 and unimodal distributions yield values *b < b*_*u*_. Conversely, distributions with *b > b*_*u*_ are either bi-modal or at least exhibit broad tails. In either case large *b*’s are a good indicator of diversification. To be conservative, the threshold for diversification is set to *b >* 0.8. (*ii*) the continuous snowdrift game with quadratic costs and benefits admits at most one single singular strategy *x*^*\**^. This implies that if *x*^*\**^ is a repellor, i.e., in the case of bi-stability, initial investments below the singular investment of the repellor, *x*_0_ *< x*^*\**^, evolve towards the investment minimum, which is always zero, while initial investments with *x*_0_ *> x*^*\**^ evolve to the investment maximum, *x*_max_ = 1. Thus, if the evolutionary outcome for *x*_0_ = 0.1 is a low mean investment 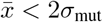, the simulation is repeated with the same parameters for *x*_0_ = 0.9. Parameter regions where the evolutionary outcomes depend on the initial condition are marked as bi-stable.

### S3 Evolutionary divergence of coexisting traits

For mutant traits *y* capable of invading a resident population *x*, i.e. if the invasion fitness *f*(*y, x*) *>* 0, ecological (frequency) dynamics drives the spread of the mutant trait, possibly replacing the resident. However, if the “resident” is equally capable of invading the “mutant”, *f*(*x, y*) *>* 0, then ecological dynamics drives the two traits to a co-existence equilibrium of two *resident* traits *x* and *y*. The direction of selection for further evolution in such populations with two (or more) co-existing traits is again determined by the selection gradients. Each of the coexisting resident traits, say *x*_1_,…, *x*_*m*_, has a corresponding selection gradient *D*_*i*_, *i* = 1,…, *m*, which is derived from the invasion fitness *f*(*y, x*_1_,…, *x*_*m*_) of a rare mutant *y* occurring in the resident *x*_*i*_:

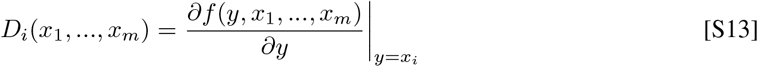

The invasion fitness of a rare mutant in trait *i* generally depends on all residents traits, and hence so do the selection gradients *D*_*i*_. Given two resident traits emerging from evolutionary diversification in a monomorphic ancestral population, the most basic question is whether the selection gradients in the two residents tend to increase the phenotypic distance between the coevolving traits, i.e., whether the selection gradients lead to evolutionary divergence.

In structured populations with two residents *x*_1_ and *x*_2_ an analytical derivation of the invasion fitness of a mutant *y, f*(*y, x*_1_, *x*_2_), is based on the local configuration of the neighbourhoods of the two residents as well as of the mutant, which together determine their respective payoffs and birthrates. Because of the time scale separation between the slow global frequency dynamics and the fast equilibration of local configurations, the neighbourhood of the two residents, *x*_1_ and *x*_2_, is approximated by the equilibria 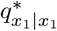 and 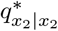 (see Eq. 4 in the main text). These local configurations determine the frequencies of *x*_1_ and *x*_2_ at their ecological equilibrium because the condition 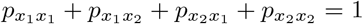 for the pair frequencies expressed in terms of 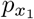 yields 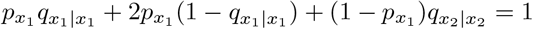 or

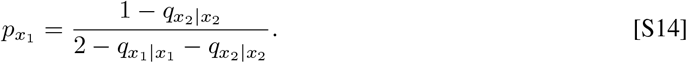

If a *x*_*i*_ resident dies, the neighbourhood of a mutant neighbour *y*_*j*_ = *x*_*j*_ + Δ, where Δ *«* 1 denotes a small mutational change in the parental trait *x*_*j*_, includes (*i*) the demised *x*_*i*_, (*ii*) one *y*_*j*_ neighbour, as well as (*iii*) 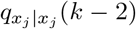 neighbours with trait *x*_*j*_ and (*iv*) 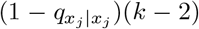 neighbours with the ‘other’ trait 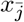, i.e. if *j* = 1 then 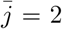 and vice versa. Note that (*ii*) follows because rare mutants *y*_*i*_ also undergo rapid local equilibration, which again yields 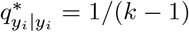 up to leading order, while (*iii*) and (*iv*) assume that the local correlations are otherwise the same as for their *x*_*j*_ parent. The resulting birthrate of the *y*_*j*_ neighbour is given by

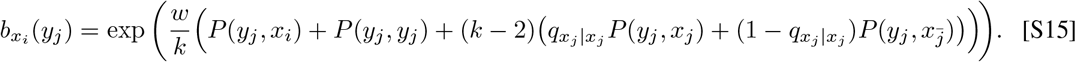

If the birthrate of *y*_*j*_ exceeds that of *x*_*j*_ it eventually takes over as a new resident replacing *x*_*j*_. Thus, divergence focusses on fitness differences between *y*_*j*_ and *x*_*j*_. On average, the birthrate of *y*_*i*_ is thus given by 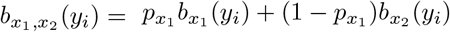 and the divergence becomes

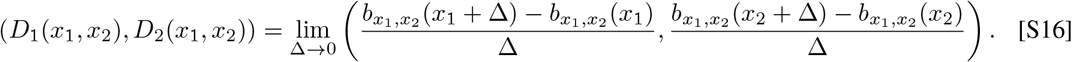

The resulting gradient field is depicted in the mutual invasibility plots for Δ = 10^*−*3^ in Fig. 5 in the main text.

### S4 Spatial adaptive dynamics: birth-death updating

For structured populations with birth-death updating, a parent is first selected from the entire population with a probability proportional to its birthrate, and then its offspring replaces one of the parent’s *k* neighbours, selected uniformly at random. Thus, with birth-death updating, competition occurs at the scale of the entire population, rather than just locally. Neighbours of individuals with high birthrates tend to be short-lived, whereas those with neighbours having low birthrates tend to live longer. This results in non-uniform life expectancies, with high turn-over in high payoff regions and low turn-over in low payoff regions. For strong selection the low birthrates can result in almost frozen regions. In general, because of global competition, outcomes of birth-death processes are more similar to outcomes in well-mixed populations than are outcomes of the corresponding death-birth processes (always assuming that selection acts on birth rates).

The setup for pair approximation with birth-death updating is basically the same as in SI Text S1: the frequency of mutants and mutant pairs increase whenever a mutant reproduces and its offspring replaces a resident neighbour. For a mutant with *j* mutant neighbours this happens with probability:

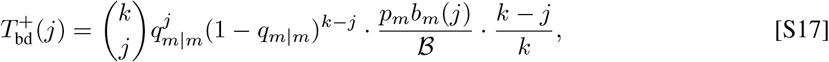

where *B* = *p*_*m*_*b*_*m*_(*kq*_*m*|*m*_) + (1 *− p*_*m*_)*b*_*r*_(*kq*_*m*|*r*_) represents the average birth rate in the population (recall that *q*_*m*|*r*_ = *p*_*m*_(1 *− q*_*m*|*m*_)*/*(1 *− p*_*m*_)), and *b*_*r*_(*i*), *b*_*m*_(*i*) denote the birth rates of residents and mutants with an (expected) number of *i* mutant neighbours. The first term in Eq. S17 represents the probability that the mutant parent has *j* mutant neighbours, the second term indicates the probability that this mutant is selected for reproduction and the last term the probability that one of its *k − j* resident neighbours is replaced. Similarly, the frequencies of mutants and mutant pairs decrease if a resident is selected for reproduction and replaces a mutant. For a resident with *j* mutant neighbours this happens with probability:

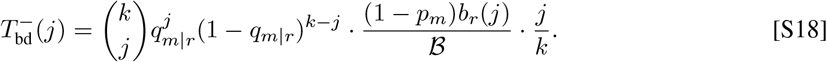

All other transitions do not alter the composition of the population. Together 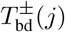 define the rate of change of the frequency of mutants:

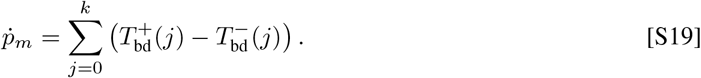

In order to derive the rates of change of *q*_*m*|*m*_ note that mutant pairs change at a rate proportional to *±j* because the parent has *j* mutant neighbours. Thus,

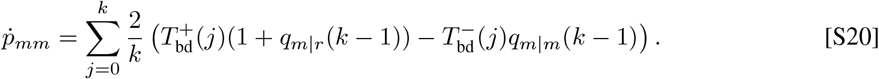

Again, the term 1*/N* in Eqs. S19 and S20 has been absorbed through a constant rescaling of time. Similarly, both equations share the common factor 1*/B*, which can be absorbed through a non-linear rescaling of time (because *B >* 0). Neither scaling changes the direction of selection or the location of equilibrium points. Note that the summations in Eqs. S19 and S20 can be carried out. Finally, using 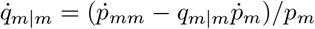 and *q*_*m|r*_ = (1 −; *p*_*m*_/(1 − *p*_*m*_) we obtain

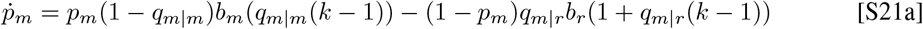

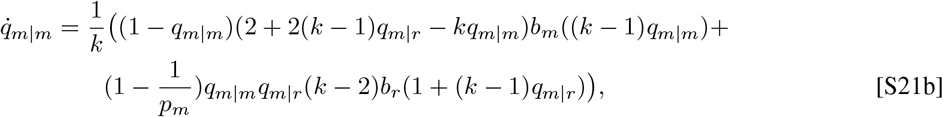

The invasion fitness of mutants, *f*(*y, x*), is the per capita growth rate, 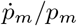, in the limit *p*_*m*_ *→* 0:

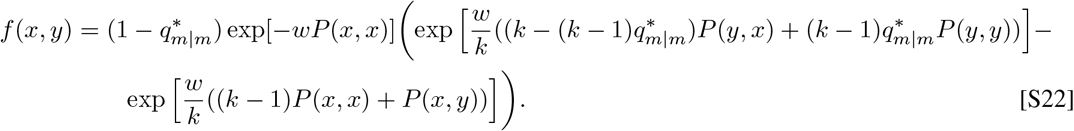

Here 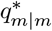 is the solution of Eq. S21b in the limit *p*_*m*_ *→* 0: for *p*_*m*_ *«* 1 the time scales of the local (fast) and global (slow) dynamics again separate (see Eq. S21), such that the local densities of mutants can be assumed to be at equilibrium 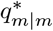. In the limit *p*_*m*_ *→* 0 the local dynamics, Eq. S21b, reduces to

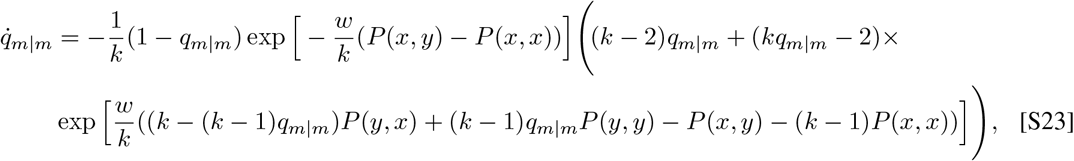

but analytical solutions remain inaccessible. However, since adaptive dynamics is based on the assumption that differences between residents and mutants are small, |*x − y*| *«* 1, we consider a Taylor expansion in *y* around *x* of the right-hand-side of Eq. S23 to obtain the first order approximation of 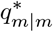:

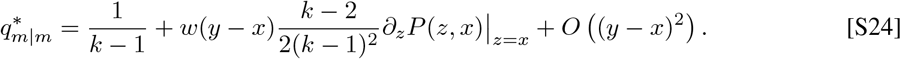

Note that the zeroth order approximation is the same as for death-birth updating (c.f. Eq. 4 in the main text). The selection gradient for birth-death updating, *D*_bd_(*x*), is now obtained by inserting Eq. S24 into Eq. S22 and evaluating at *y* = *x*:

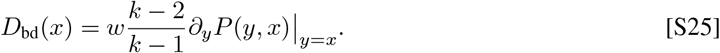

In particular, as opposed to death-birth updating (c.f. Eq. 5 in the main text) spatial structure does not affect the sign of the selection gradient as compared to well-mixed populations (Doebeli et al., 2004) and hence spatial structure does not affect the existence or location of singular points *x*^*\**^. The Jacobian of the selection gradient and the Hessian of the fitness at *x*^*\**^ are:

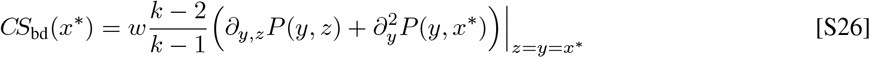

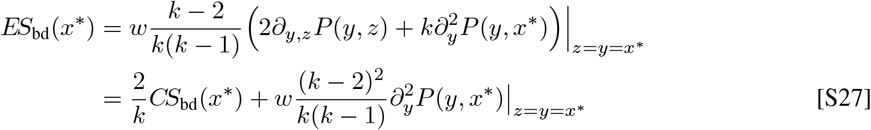

The conditions for convergent stability are again the same as in well-mixed populations, whereas the criteria for evolutionary stability are generally different.

#### S4.1 Continuous prisoner’s dilemma

In the continuous prisoner’s dilemma, the selection gradient for the spatial adaptive dynamics is

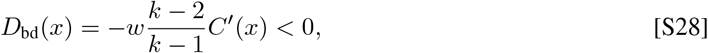

such that for increasing cost functions the selection gradient is always negative and investment levels invariably evolve to zero – just as in well-mixed populations. Consequently, no singular strategy exists and it is impossible to maintain cooperation no matter how significant the benefits of cooperation may be. Accordingly, for the linear continuous prisoner’s dilemma investment levels consistently hover near zero in individual-based simulations, regardless of the cost-to-benefit ratio *r*. Evolutionary trajectories are essentially the same for linear or non-linear variants of the prisoner’s dilemma and indistinguishable from the scenario depicted in Fig. 2d-f in the main text, where defection prevails.

#### S4.2 Continuous snowdrift game

In the continuous snowdrift game the selection gradient for spatial adaptive dynamics with birth-death updating, Eq. S25, becomes:

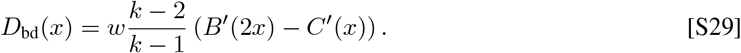

Similarly, convergence and evolutionary stability of a singular strategy *x*^*\**^ are given by the conditions

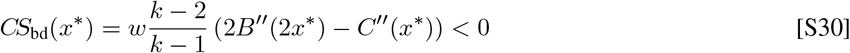

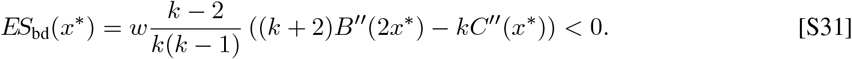

The condition for convergence stability is again the same as in well-mixed populations, while evolutionary stability differs (c.f. Eqs. 6, 7 in the main text and (Doebeli et al., 2004)). For quadratic cost and benefit functions the condition for evolutionary stability is *b*_2_(*k* + 2) *< c*_2_*k* and converges to that of well-mixed populations (*b*_2_ *< c*_2_) for large *k*. Combining the two stability conditions shows that evolutionary branching occurs for 2*b*_2_ *< c*_2_ *< b*_2_(*k* + 2)*/k*, which is more restrictive than in the well-mixed game, but not as significant as for spatial structures with death-birth updating (c.f. Fig. S1a and Fig. 4a in the main text). However, systematic deviations from the analytical predictions can be observed in individual-based simulations resulting in an increase of the parameter range generating evolutionary diversification (see Fig. S1). In contrast to death-birth updating, spatial structure promotes diversification for weak selection to a greater extent (c.f. Fig. S1b and Fig. 4d in the main text). More specifically, evolutionary diversification extends into regions where pair approximation predicts (*i*) full investments and (*ii*) intermediate ESS’s. For birth-death updating the turn-over in areas with low birthrates can be very slow and result in (almost) frozen regions. However, those frozen traits can become advantageous again at a later point in time and contribute to the process of diversification. As a result, this can introduce historical contingencies where the evolutionary end state depends on the initial configuration. Otherwise, the reasons for the discrepancies between analytical approximations and individual-based simulations are essentially the same as in structured populations with death-birth updating: finite-size mutations and positive population variance allow the individual-based process to reach regions in traits space that are prohibited in the limit of infinitely small mutations, and allow for coexistence and evolutionary divergence between coexisting strains.

**Fig. S1.**
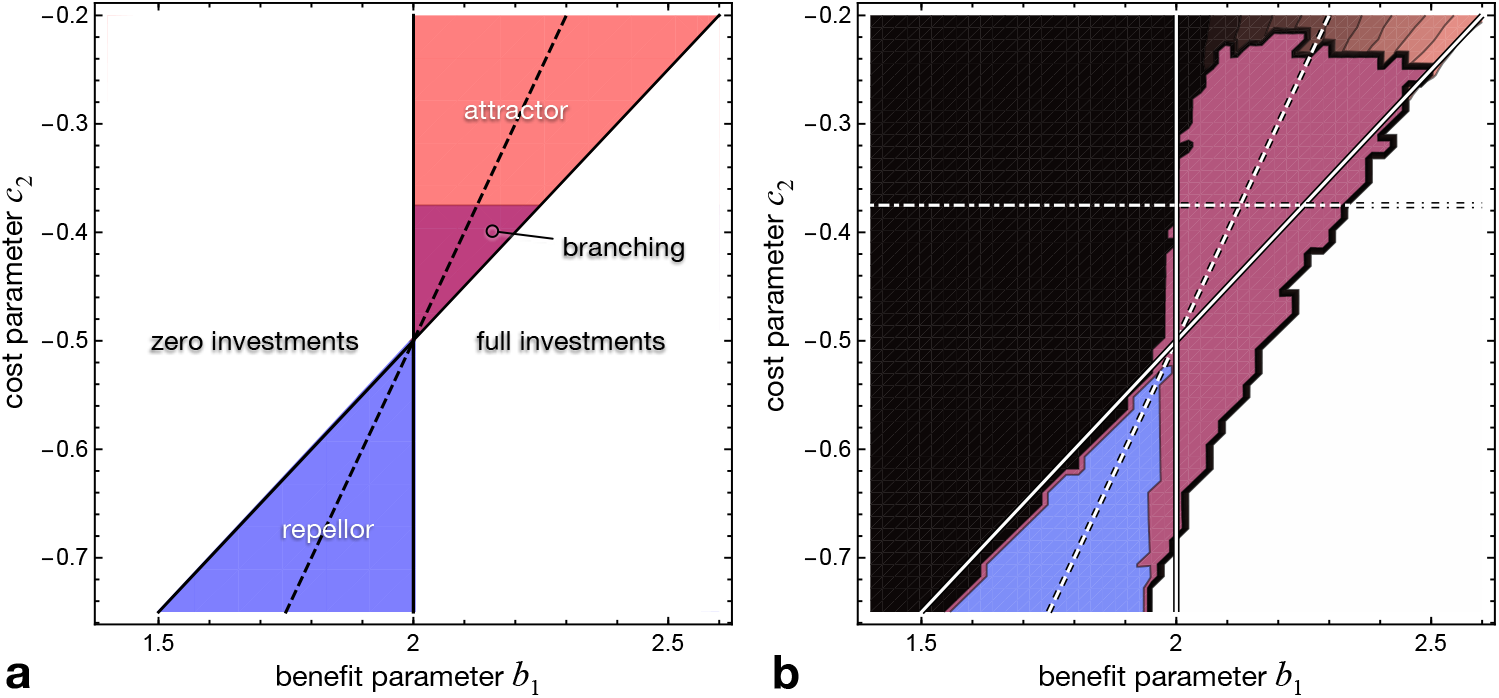
Continuous snowdrift game with quadratic benefit and cost functions, *B*(*x*) = *b*_1_*x* + *b*_2_*x*^2^ and *C*(*x*) = *c*_1_*x* + *c*_2_*x*^2^. Evolutionary outcomes are shown as a function of the benefit parameter *b*_1_ and cost parameter *c*_2_ with *b*_2_ = *−*1*/*4 and *c*_1_ = 2. Note that for *b*_1_ *<* 2 *B*(*x*) *< C*(*x*) holds at least for small *x*, which means that lower investors invariably dominate and effectively recover the characteristics of the prisoner’s dilemma. The predictions by adaptive dynamics **a** differ from well-mixed populations only in that the parameter region of evolutionary branching is smaller (c.f. Fig. 4a in the main text) while all other thresholds are unchanged. In contrast, individual-based simulations **b**, where equilibrium investment levels range from the minimum (black) to intermediate (grey) and the maximum (white) augmented by convergent stability (red), bi-stability (blue), and diversification (maroon), indicate much larger regions of parameter space that result in diversification than predicted by evolutionary branching (click here for interactive online simulations).

## References

B. Allen, M. A. Nowak, and U. Dieckmann. Adaptive dynamics with interaction structure. American Naturalist,181(6):E139–E163, 2013. ISSN 00030147.

B. Allen, G. Lippner, Y.-T. Chen, B. Fotouhi, N. Momeni, S.-T. Yau, and M. A. Nowak. Evolutionary dynamics on any population structure. Nature,544:227–230, 2017.

R. M. Dawes. Social dilemmas. Annual Review of Psychology,31:169–193, 1980.

F. Débarre, C. Hauert, and M. Doebeli. Social evolution in structured populations. Nature Communications, 5 (3409), 2014.

U. Dieckmann and R. Law. The dynamical theory of coevolution: a derivation from stochastic ecological processes. Journal of Mathematical Biology,34:579–612, 1996.

M. Doebeli. Adaptive Diversification. Monographs in Population Biology. Princeton University Press, Princeton, NJ, 2011.

M. Doebeli and C. Hauert. Models of cooperation based on the prisoner’s dilemma and the snowdrift game. Ecology Letters,8:748–766, 2005.

M. Doebeli, C. Hauert, and T. Killingback. The evolutionary origin of cooperators and defectors. Science, 306 (5697):859–62, Oct 2004.

M. Doebeli, C. Hauert, and T. Killingback. A comment on “towards a rigorous framework for studying 2-player continuous games” by shade t. shutters, journal of theoretical biology 321, 40–43, 2013. Journal of Theoretical Biology,336:240–241, 2013.

J. B. Freeman and R. Dale. Assessing bimodality to detect the presence of a dual cognitive process. Behavior Research Methods,45:83–97, 2013.

F. Fu, M. A. Nowak, and C. Hauert. Invasion and expansion of cooperators in lattice populations prisoner’s dilemma vs. snowdrift games. Journal of Theoretical Biology,266(3):358–366, Jul 2010.

S. A. H. Geritz, J. A. J. Metz, É. Kisdi, and G. Meszéna. Dynamics of adaptation and evolutionary branching. Physical Review Letters,78(10):2024–2027, Feb 1997.

S. A. H. Geritz, E. Kisdi, G. Meszéna, and J. A. J. Metz. Evolutionarily singular strategies and the adaptive growth and branching of the evolutionary tree. Evolutionary Ecology Research,12:35–57, 1998.

D. Greig and M. Travisano. The Prisoner’s Dilemma and polymorphism in yeast SUC genes. Biology Letters,271:S25 – S26, 2004.

G. Hardin. The tragedy of the commons. Science,162:1243–1248, 1968.

C. Hauert. EvoLudo: Interactive tutorials on evolutionary game theory, https://www.evoludo.org, 2021.

C. Hauert and M. Doebeli. Spatial structure often inhibits the evolution of cooperation in the snowdrift game. Nature,428:643–646, 2004.

C. Hauert, F. Michor, M. A. Nowak, and M. Doebeli. Synergy and discounting of cooperation in social dilemmas. Journal of Theoretical Biology,239:195–202, 2006.

R. Ibsen-Jensen, K. Chatterjee, and M. A. Nowak. Computational complexity of ecological and evolutionary spatial dynamics. Proceedings of the National Academy of Sciences USA,112(51):15636–15641, 2015.

T. Killingback, M. Doebeli, and N. Knowlton. Variable investment, the continuous prisoner’s dilemma, and the origin of cooperation. Proceedings of the Royal Society B,266:1723–1728, 1999.

T. Killingback, M. Doebeli, and C. Hauert. Diversity of cooperation in the Tragedy of the Commons. Biological Theory,5(1):3–6, 2010.

P. Langer, M. A. Nowak, and C. Hauert. Spatial invasion of cooperation. Journal of Theoretical Biology,250: 634–641, 2008.

J.-F. Le Gaillard, R. Ferriére, and U. Dieckmann. The adaptive dynamics of altruism in spatially heterogenous populations. Evolution,57:1–17, 2003.

H. Matsuda, N. Ogita, A. Sasaki, and K. Sato. Statistical mechanics of populations. Progress of Theoretical Physics,88(6):1035–1049, 1992.

J. A. J. Metz, S. A. H. Geritz, G. Meszena, F. J. A. Jacobs, and J. S. van Heerwaarden. Adaptive dynamics: a geometrical study of the consequences of nearly faithful replication. In S. J. van Strien and S.M. Verduyn Lunel, editors, Stochastic and Spatial Structures of Dynamical Systems, pages 183–231. North Holland, Amsterdam, 1996.

M. A. Nowak and R. M. May. Evolutionary games and spatial chaos. Nature,359:826–829, 1992.

H. Ohtsuki, C. Hauert, E. Lieberman, and M. A. Nowak. A simple rule for the evolution of cooperation on graphs. Nature,441:502–505, 2006.

K. Parvinen, H. Ohtsuki, and J. Y. Wakano. The effect of fecundity derivatives on the condition of evolutionary branching in spatial models. Journal of Theoretical Biology,416:129–143, 2017.

K. Sigmund. The Calculus of Selfishness. Princeton Univ. Press, 2010.

R. Sugden. The economics of rights, co-operation and welfare. Blackwell, Oxford and New York, 1986.

G. Szabó and G. Fáth. Evolutionary games on graphs. Physics Reports,446:97–216, 2007.

G. Szabó and C. Hauert. Phase transitions and volunteering in spatial public goods games. Physical Review Letters,89:118101, 2002.

C. E. Tarnita, N. Wage, and M. A. Nowak. Multiple strategies in structured populations. Proceedings of the National Academy of Sciences USA,108:2334–2337, 2011.

M. van Baalen. Pair approximations for different spatial geometries. In U. Dieckmann, R. Law, and J. A. J. Metz, editors, The geometry of ecological interactions: Simplifying spatial complexity, Cambridge Studies in Adaptive Dynamics,pages 359–389. Cambridge University Press,Cambridge, 2000.

